# Muscarinic modulation of spike-timing dependent plasticity at recurrent layer 2/3 synapses in mouse auditory cortex

**DOI:** 10.1101/690446

**Authors:** Deepti Rao, Megan B. Kratz, Paul B. Manis

**Affiliations:** Department of Cell Biology and Physiology, University of North Carolina at Chapel Hill, Chapel Hill, NC; Department of Otolaryngology/Head and Neck Surgery, University of North Carolina at Chapel Hill, Chapel Hill, NC

**Keywords:** calcium, back-propagation, acetylcholine, long-term potentiation, long-term depression, NMDA receptors

## Abstract

Cholinergic systems contribute to the refinement of auditory cortical receptive fields by activating muscarinic acetylcholine receptors (mAChRs). However, the specific cellular and synaptic mechanisms underlying acetylcholine’s effects on cortical circuits are not fully understood. In this study, we investigate the effects of muscarinic receptor modulation on spike-timing dependent plasticity (STDP) at synapses onto layer 2/3 pyramidal neurons in mouse auditory cortex (AC). Synapses onto layer 2/3 pyramidal neurons exhibit a STDP rule for pairing of postsynaptic spike bursts with single presynaptic stimuli. Pre-before-post pairing at +10 ms results in a timing-dependent long-term potentiation (tLTP), whereas pre-before-post pairing at +50 ms intervals, and post-before-pre pairing at -10 to -20 ms produce a timing-dependent long-term depression. We also characterize how mAChR activation affects plasticity at these synapses, focusing on the induction of tLTP. During pre-before-post pairing at +10 ms, mAChR activation by either carbachol or oxotremorine-M suppresses tLTP. mAChR activation also reduces the NMDA-receptor dependent synaptically evoked increase in calcium in dendrites, apparently without affecting presynaptic transmitter release. Pharmacological experiments suggest that M1 and M3 receptors are not involved in the mAChR-mediated suppression of tLTP. Taken together, these results suggest activating mAChRs in layer 2/3 intracortical circuits can modify the circuit dynamics of AC by depressing tLTP mediated by NMDA receptors, and depressing calcium influx at excitatory synapses onto layer 2/3 pyramidal cells.

## Introduction

Experience-dependent plasticity contributes to organizing the representation and processing mechanisms of sensory information in auditory, visual and somatosensory cortices (Allen et al. 2003; Chang and Merzenich 2003; Hubel and Wiesel 1965; Kotak et al. 2007; Rao et al. 2010; Takesian et al. 2012; de Villers-Sidani et al. 2007; Xu et al. 2007). Information about the context and behavioral significance of sensory information is provided in part by the activity of neuromodulatory systems. In the auditory system, these systems are critical for shaping experience-dependent plasticity of sensory representations (Kilgard 1998; Metherate et al. 2005; Metherate and Weinberger 1989; Weinberger 2015; Zhang et al. 2005). At a cellular level representational plasticity is hypothesized to depend on correlations between pre and postsynaptic activity, and to require long-term potentiation (LTP) and depression (LTD) of synapses (Buonomano and Merzenich 1998). Although there are many demonstrations that neuromodulation can engage or prevent representational plasticity in neocortex, the mechanisms by which changes to synaptic strength are regulated by specific modulators are not well understood.

In the primary auditory cortex (A1), intracortical and thalamic inputs combine to determine the sensory responses of individual neurons (Intskirveli et al. 2016; Kaur et al. 2005; Liu et al. 2007; Winkowski and Kanold 2013). Part of the intracortical circuit in layer 2/3 is formed by the horizontal axons of layer 2/3 pyramidal cells, which can link columns of neurons with different frequency tuning (Clarke et al. 1993; Matsubara and Phillips 1988; Ojima et al. 1991; Read et al. 2002; Song et al. 2006; Watkins et al. 2014). Physiological studies have shown that layer 2/3 pyramidal neurons have broad sub-threshold receptive fields (Kaur et al. 2004; Liu et al. 2007; Ojima 2002), and receive inputs tuned to a wide range of frequencies, even on adjacent spines (Chen et al. 2011). The convergence of inputs across the tonotopic map could play an important role in integrating responses to spectrotemporally complex stimuli commonly encountered in the acoustic environment, such as frequency modulated sounds or sounds with harmonic structures (Harper et al. 2016; Kadia and Wang 2003; Kratz and Manis 2015), and can create a substrate for a flexible activity-dependent plasticity of suprathreshold sensory responses and neurons sensitive to multiple acoustic features (Atencio and Sharpee 2017; Harper et al. 2016).

Plasticity of sensory representations has been associated with spike timing-dependent plasticity (STDP) in vivo in auditory, visual and somatosensory cortices (D’amour and Froemke 2015; Dahmen et al. 2008; Jacob et al. 2007; Larsen et al. 2010; Martins and Froemke 2015; Yao and Dan 2001). STDP involves changes in strength of synapses that is dependent upon the precise timing of pre- and postsynaptic activity (Bi and Poo 1999; Markram 1997). Most commonly, presynaptic activity that precedes postsynaptic firing by tens of milliseconds can increase synaptic strength (timing-dependent LTP; tLTP), whereas reversing this temporal order can weaken synaptic strength (timing-dependent LTD; tLTD). tLTP largely relies on the interplay between NMDA receptor activation and the timing of back-propagating action potentials in the dendrites of the postsynaptic neuron (Linden 1999; Magee and Johnston 1997; Sourdet 1999), whereas tLTD can result either from postsynaptic NMDA receptor activation (Karmarkar et al. 2002) or from a cascade involving postsynaptic metabotropic glutamate receptors and presynaptic cannabinoid receptors (Bender 2006; Nevian and Sakmann 2006). The temporal shape and direction of the STDP window varies with brain region, cell and synapse type (Larsen et al. 2010).

Activation of cholinergic receptors has been widely implicated in the modulation of tonotopic map plasticity in auditory cortex. Pairing acoustic stimuli with electrical stimulation of nucleus basalis, which provides cholinergic innervation of the cortex, alters the subsequent representation of the stimulus (Froemke et al. 2007; Kilgard 1998; Weinberger 2003). Muscarinic cholinergic receptors (mAChRs) have been shown to modulate sensory responses as well as transmission at both excitatory and inhibitory synapses in auditory cortex (Atzori et al. 2005; Bajo et al. 2014; Flores-Hernandez et al. 2009; Kuchibhotla et al. 2017; Metherate and Ashe 1991, 1995; Metherate and Weinberger 1990). mAChRs also play a crucial role in the development and function of the normal auditory cortex. Absence of mAChRs leads to a distorted tonotopic map and a decrease in tonotopic map plasticity (Zhang et al. 2006). Attenuation of cholinergic inputs to cortex disrupts auditory spatial perception and plasticity (Leach et al. 2013) and performance of a learned task when performing in an active, but not passive, context (Kuchibhotla et al. 2017). Even though the cholinergic system plays an important role in auditory cortex, it remains unclear how acetylcholine influences tLTP and tLTD at cortical synapses, and which classes of receptors mediate specific effects.

In this study, we investigated the timing rules of STDP and their modulation by one set of receptors activated by cholinergic system, the muscarinic receptors, at recurrent synapses in layer 2/3 of the mouse auditory cortex. We find that the STDP in auditory cortex follows unique timing rules, in which tLTP occurs at +10 ms (presynaptic EPSP leading postsynaptic spikes), while tLTD occurs at both -10 and +50 ms. Activation of mAChRs modulates the timing rules. The muscarinic agonists carbachol and oxotremorine-M gate tLTP induction at +10 ms and also decrease NMDA receptor currents in layer 2/3 cells. The tLTP depends on increases in intracellular calcium, and during pairing of presynaptic and postsynaptic activity with a +10 ms delay, carbachol application results in a decrease in action potential evoked postsynaptic calcium influx, as well as a decrease in the summed action potential and synaptic calcium influx, an effect that is likely caused by the reduction in NMDAR currents. We conclude that layer 2/3 synapses in auditory cortex exhibit STDP, and that this plasticity can be modulated through a postsynaptic mechanism by activation of muscarinic cholinergic receptors.

## Materials and Methods

The experiments reported here were performed in two groups. The first set of experiments was performed from 2008-2011, and a second set of experiments was performed from 2015-2016. All protocols were performed according to methods approved by the Institutional Animal Care and Use Committee of the University of North Carolina, Chapel Hill. Thalamocortical brain slices of the auditory cortex were prepared from young CBA/CaJ mice (P10–P22; most were in the range P13-17;), following procedures previously described (Cruikshank et al. 2002; Kratz and Manis 2015; Rao et al. 2010). Mice of either sex were used in both the first and second set of experiments. Sex was undetermined except in a small subset of mice used in the second set of experiments, and those sex determinations are indicated in the text.

To prepare brain slices, mice were first anesthetized with 100 mg/kg ketamine, 10 mg/kg xylazine, and decapitated when areflexic. The brain was trimmed, and slices were cut in a chilled solution at an angle of +15 degrees from the horizontal plane, so as to include A1 and key landmarks of the thalamocortical pathway. The rostral-caudal axis of these slices is approximately parallel to the tonotopic axis of the primary auditory region. Because of variability in the organization of the auditory cortical map in mouse (Issa et al. 2014; Tsukano et al. 2015), no attempt was made to limit recordings from a given tonotopic area; thus data are pooled from recordings made in both “primary” and “shell” slices, following the prior description (Cruikshank et al. 2002). Consequently, the recording sites are referred to here as auditory cortex (AC), and we reserve the use of the term A1 to specifically reference prior *in vivo* work where precise knowledge of the cortical fields is available. The standard slicing, incubation, and recording solution was an artificial cerebrospinal fluid (ACSF) that contained (in mM) 134 NaCl, 3.0 KCl, 2.5 CaCl_2_, 1.3 MgCl_2_, 1.25 KH_2_PO_4_, 10 glucose, 20 NaHCO_3_, 0.4 ascorbic acid, 2 sodium pyruvate, and 3 myoinositol; this solution was saturated with 95%O_2_/5%CO_2_. In the early experiments, the slices were incubated at 34°C during a 1-hour recovery period, whereas in the later experiments, slices were incubated at 34°C for a 15-minute period. In both sets of experiments, slices were subsequently maintained at room temperature until used (> 1 hr after slice preparation).

### Recording and stimulation

The recording and stimulating arrangements are illustrated in Figure 1. During recording, single cells were visualized with infrared interference contrast optics or asymmetric illumination, and recorded using patch pipettes in current-clamp, or in a subset of experiments investigating NMDA receptor function, in voltage-clamp. A concentric bipolar stimulating electrode (Fredrick Haer, #CBBB75, 75µM diameter) was placed in layer 2/3 in AC, 500-700 µm caudal or rostral to the recording site (Figure 1A). EPSPs were evoked by computer-generated monophasic 100 µsec pulses delivered through an optically isolated analog stimulator (Dagan S940 and S910 isolation unit) through the stimulating electrode. Cortical pyramidal cells in auditory cortex layer 2/3 were identified on the basis of their position (Figure 1A) and electrophysiological (Figure 1B) criteria (Rao et al. 2010). Fast-spiking cells (interneurons) were identified based on spike width and firing rate, and were excluded in this study. All recordings were performed using whole-cell tight seal methods at 34°C.

**Figure 1:**
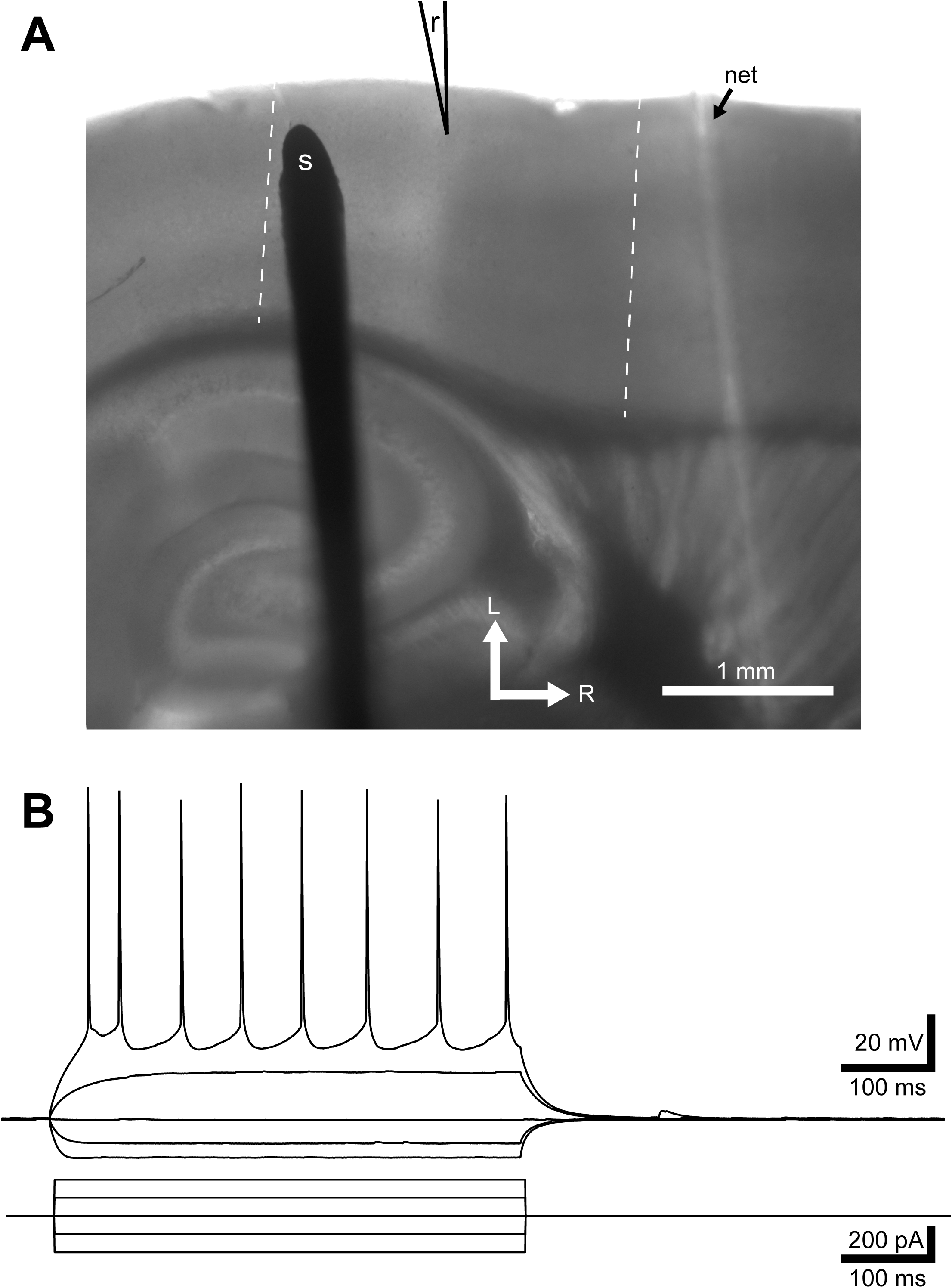
Arrangement of stimulating and recording electrodes in layer 2/3 of auditory cortex. A. Low-magnification view of a thalamocortical slice showing the location of the stimulating electrode (*s*) and the recording electrode (*r*) during an experiment. The boundaries of the cortical area examined in this study are indicated by the dashed white lines. An indentation associated with the net holding the slice in the chamber is also indicated (*net*). B. Current-clamp recordings in response to current pulses from a layer 2/3 pyramidal cell, showing the regular firing with adaptation typical of the recorded cells in this study. Top: Voltage traces. Bottom: Injected current steps.

### Current Clamp Recordings and STDP Protocols

Recording pipettes were pulled from 1.5 mm KG-33 glass and backfilled with an intracellular solution containing (in mM) 130 K-gluconate, 4 NaCl, 0.2 EGTA, 10 HEPES, 2 Mg_2_ATP, 0.3 Tris GTP, and 10 phosphocreatine (pH 7.2 with KOH), and 290 mOsm. In a few experiments in the second group, the pipette Mg_2_ATP was 4 mM; no differences in STDP or cell excitability were evident however. In one set of experiments, the electrode solution was supplemented with 10 mM BAPTA. Cells were first briefly characterized with a current-voltage pulse protocol to confirm firing patterns and cell health. Next, thresholds for action potential generation were evaluated for 2-5 ms current pulses. Baseline excitatory postsynaptic potentials (EPSPs) were evoked every 10 seconds by stimulating in layer 2/3 with 100 µsec shocks to activate presynaptic fibers. The amplitudes of the EPSPs were targeted to be 5-8 mV in the first set of experiments, and 2-3 mV in the second set of experiments. During the STDP induction protocols, the postsynaptic cell was also stimulated with a 5-pulse train at 125 Hz of 1-3 nA depolarizing current pulses, 2-5 ms in duration, to generate a train of 5 action potentials with a fixed delay with respect to the presynaptic stimulation. The high-frequency burst was used to maximize the generation of an AP-induced calcium influx over a wide area of the dendritic tree. The induction protocol consisted of 100 presentations of the EPSP-AP burst combination, at an interval of 1/s. During the induction protocol for pre-before-post pairing, the spike-EPSP was measured from the onset of the evoked EPSP to the peak of the first postsynaptic action potential. For post-before-pre pairing, the timing was measured from the peak of the 5th action potential to the onset of the EPSP (Nevian and Sakmann 2006). The timing is indicated in the insets of Figure 2.

**Figure 2:**
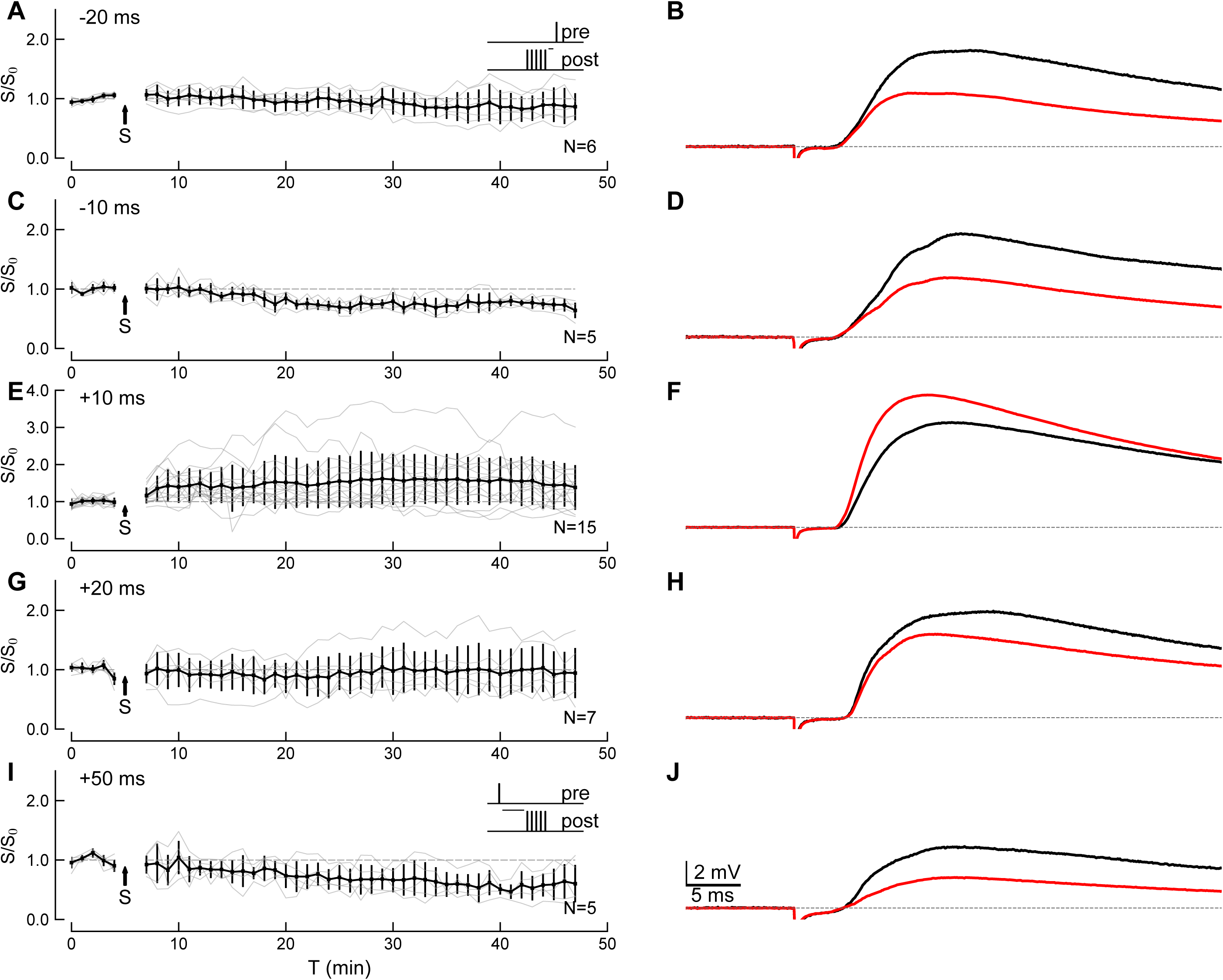
Changes in the strength of layer 2/3 auditory cortical synapses onto pyramidal cells induced by repetitive pairing of EPSPs with postsynaptic spike bursts. For each pairing interval, the left column summarizes the time course of the maximum EPSP onset slope relative to the baseline slope (*S/S_0_*). The right column shows example EPSPs before and after pairing. A. −20 ms (post→pre) leads to weak tLTD (average control EPSP is shown in black, and the averaged post-pairing EPSP is shown in red). The inset in A shows the pairing paradigm for negative intervals (panels A-D), where the EPSP is elicited by presynaptic fiber stimulation, and the postsynaptic cell is forced to spike with a train of 5 current pulses at 8 ms intervals. The pairing occurs at the time indicated by *S* and the arrow in the graph in this and subsequent figures. The time indicated by the short horizontal line in the inset represents the measurement of the pairing interval. *N* indicates the number of cells. B. Comparison of averaged EPSPs during the baseline (black trace) and over 25-40 minutes after pairing (red trace). The same conventions are used in the remaining panels. C, D. −10 ms post→pre pairing results in tLTD. E, F. Summary of pre→post pairing at +10 ms, with EPSPs preceding the postsynaptic spikes, results in tLTP. G, H. Same paradigm as E, with pre→post pairing at +20 ms showing no change on average. I, J. Summary for pre→post pairing at +50 ms, resulting in tLTD. The inset in I illustrates the pairing paradigm used in panels E-J. The time indicated by the horizontal line in the inset corresponds to the pairing interval. The time course of EPSP slope for individual cells is shown by the faint gray lines. Error bars are SDs.

Following the induction protocol, EPSPs were monitored every 10 seconds for the next 30-40 minutes. EPSPs frequently had a compound appearance with inflections on both the rising and falling phases, suggesting the activation of multiple inputs or polysynaptic pathways. To focus on synaptic inputs that were most likely to be monosynaptic and not contaminated with polysynaptic events, the maximal slope of the first 2-3 ms of the EPSP was measured. The EPSP slopes were averaged in 1-minute blocks (6 sequential trials). The EPSP slope ratio (S/S_0_) for each cell was then computed as the ratio of the mean EPSP slope 20–40 min after the induction protocol (S) to the mean slope measured during the 5-min baseline prior to induction protocol (S_0_). Cells were retained for analysis if they had resting membrane potentials negative to -60 mV, exhibited regular firing, showed less than an 8-mV shift in resting membrane potential during the protocol, and were stable for at least 30 minutes after the induction protocol. Fast-spiking cells, and one cell with a very high adaptation ratio that was tested with a +10 ms interval in eserine were excluded from further analysis. In addition, cells that showed a coefficient of variation of EPSP slope (measured as the standard deviation/mean of the 1-minute averaged EPSPs) > 0.30 during the baseline period were excluded when analyzing the STDP data.

To assess the effect of mAChR activation in these experiments, a cholinergic agonist (20 μM carbachol or 3 μM oxotremorine-M) was applied. In the first set of experiments, a computer-controlled valve delivered the agonist to the bath from 2 minutes before to 1 minute after the onset of the induction protocol, for a total duration of 5 minutes. In the second set of experiments, a manually switched valve began delivery 3 minutes before, and returned to ACSF immediately after the induction protocol, for a total agonist delivery duration of 5 minutes. In these pharmacological experiments, the baseline was limited to the first 3 minutes, to prevent agonist application from interfering with the baseline measure. Because the solutions took ∼1 minute to reach the chamber, and ∼1 minute to fully exchange with the solution in the chamber, the slice was exposed to the agonists from just before the beginning until the end of the induction protocol with the agonist at full concentration in both sets of experiments, and the agonist was washed out of the slice afterwards. When antagonists were used, they were present in the solution for the entire duration of the recording, including while agonists were applied.

The effects of mAChR activation on basal synaptic transmission and cell excitability were measured in separate experiments, without an STDP induction protocol. In these experiments, single shocks to layer 2/3 were delivered at 0.1 Hz, and solution exchange was performed as for the STDP measurements. The cell’s firing rate versus injected current (FI) relationship was measured immediately before the baseline EPSP measurements were taken, and repeated at the end of the baseline period, immediately after the agonist delivery was discontinued, and at the end of the recording period. FI curves took 2 minutes to acquire.

FI curves were measured using a series of 500-ms duration current pulses with 20-50 pA steps up to a maximum of 200-400 pA. The firing frequency (F) at each current level (I), was parameterized by fitting the curve up to the maximum firing rate (thus excluding traces with depolarization block) to the following function:

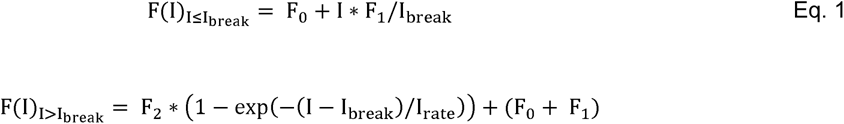

Here, F(I) is the firing rate (in spikes per second) measured with a current step of amplitude I (in pA), F_0_ is the firing rate in the absence of current (spikes per second), I_break_ (in pA) defines the breakpoint that separates the near-threshold linear region of the FI curve from the exponentially rising portion and is the threshold current for firing, F_1_ is the firing rate at I_break_, F_2_ is the increase in firing rates when the current exceeds I_break_, and I_rate_ is the rate of change during the exponentially increasing phase of firing with current level (units of pA). Fitting used the sequential least squares programming method (SQSLP) from the Python library scipy.optimize (version 0.15.1, www.scipy.org). Eq. 1 parameterizes key measures of the FI curve, including the threshold for firing, the rate at which the firing rate changes with current (the slope), and the maximum firing rate, which facilitates comparison between experimental treatments. For the fits here, we held F_0_ and F_1_ at 0. Fits were compared for the FI curves taken immediately before drug application, and immediately after drug application. The final FI curves after washout of the drugs were not analyzed because activation of mAChRs can result in long-term activation of protein kinases and phosphatases, producing effects that do not completely wash out.

### Voltage clamp recordings

In a subset of experiments, cells were voltage-clamped to isolate and measure NMDA receptor-dependent currents. For these experiments, the slices were perfused with a modified oxygenated ACSF (as above) with the following changes. Calcium and magnesium were increased to 4 mM each, AMPA receptors were blocked with 10 μM CNQX, and GABA receptors were blocked with 50 μM picrotoxin. These conditions pharmacologically isolate the NMDAR-mediated currents while minimizing polysynaptic transmission (Philpot et al. 2001a, 2001b). Recording pipettes were pulled from 1.5 mm KG33 glass, and the tips were coated with Sylgaard 184 (Dow Corning) to reduce capacitance. Electrodes were filled with an internal solution containing (in mM): 120 cesium methanesulfonate, 8 TEA-chloride, 10 HEPES, 0.2 EGTA, 4 tris-adenosine triphosphate, 0.3 tris-guanosine triphosphate, 10 creatinine phosphate, and 3 QX-314 chloride with pH adjusted to 7.2 and osmolarity adjusted to 300 mOsm with sucrose. The internal solution also contained 50-100 ≈M AlexaFluor 488 for post-hoc identification of pyramidal neurons. Pipette capacitive transients were minimized prior to breaking in, and after break-in whole cell capacitance and series resistance compensation was applied [Rs, mean=16.6 (SD 4.8) MΩ, compensation, mean=41.4 (SD 12.7) %]. Cells were stepped to +40 mV when measuring NMDA currents, and the difference between traces with and without stimulation in layer 2/3 was computed to isolate the synaptic current from the residual outward potassium currents. The residual potassium current was 570 (SD 124) pA (range: 375-846 pA), and the cell voltage after estimating the drop across the series resistance of the electrode was calculated to be +30.8 (SD 5.5) mV. Current traces were low-pass filtered at 2 kHz. The stimulus intensity was adjusted to evoke 100-500 pA EPSCs while the cells were depolarized. Stimuli were delivered either as single shocks, every 8 sec, or in 7 of the 13 tested cells, as pairs of shocks 50 ms apart to measure the paired-pulse ratio. Ten traces were averaged for each measurement.

### Electrophysiology Data Acquisition and Analysis

Data for experiments performed in 2008-2011 were acquired using a custom MATLAB program (R2008-R2018, The Mathworks, Natick, MA). Data for the calcium imaging experiments from that series, and for the experiments performed in 2015-2016, were acquired using the Python program ACQ4 (Campagnola et al. 2014), available at http://www.acq4.org.

Data were analyzed using MATLAB, ACQ4, Python scripts, Igor Pro (6.2 Wavemetrics, Oswego, OR) and Prism 6.0 and 7.0 (Graphpad, San Diego, CA). The analysis pipeline involved multiple steps. For the STDP and pharmacological measurements of the first data set, MATLAB scripts were used to generate tables of the 1-minute binned EPSP slope time courses that were organized and stored in Excel (Microsoft, V14.6.4) spreadsheets. Data from the second set of experiments were analyzed in ACQ4, and the resulting 1-minute binned EPSP slope time courses were saved as text files and added to the same Excel sheets. The Excel sheets were subsequently read using a Python script to combine cells and analyze groups using uniform metrics. For analysis of intrinsic excitability, the analyses used Python scripts that directly read the original (raw) data from both MATLAB and ACQ4 files, and then performed identical processing for both early and late data sets.

### Calcium Imaging

Calcium imaging was performed in cells under current clamp, using conditions that closely matched those used during the spike-timing protocol; however, we did not attempt to induce or measure plasticity. In the first set of experiments, pipettes (1.5 mM KG33) were filled with intracellular solution supplemented with the low-affinity calcium-indicator Fluo-5F (Life Technologies, 100 µM). Fluo-5F has a reported k_d_ for Ca^2+^ of ∼ 0.70 µM in 1 mM Mg^2+^ at 30°C (Woodruff et al. 2002), or 2.3 µM at 22°C (ThermoFisher product data sheet for catalog number F14221; Mg^2+^ concentration not specified). As our experiments were performed with 2 mM Mg^2+^ in the pipette, the k_d_ is likely higher than 0.70 µM. When using Fluo-5F, AlexaFluor 568 (50 µM) was included in the pipette solution to reveal neuronal morphology. In the second set of experiments, a higher affinity indicator, Asante Calcium Red (ACR, TEFLabs, Austin, TX; 100 µM) was used in a non-ratiometric 2+ 2+ mode. This indicator has a reported k_d_ for Ca of ∼0.53 µM in 1 mM Mg at 22°C (Hyrc et al. 2013), and 0.4 µM in 0 Mg2^+^ (TEFLabs product information). ACR was used in conjunction with Lucifer Yellow-CH (potassium salt) (<0.5%) for cell identification. Imaging and recording took place on a Zeiss FS-2 microscope under a 40X 0.75NA water immersion objective. Subsequent to whole-cell break-in, cells were monitored for a minimum of 15 min to allow diffusion of the dyes into the dendrites before fluorescence measurements were taken. Voltage and fluorescent signals were measured simultaneously. To generate EPSPs, an extracellular stimulation pipette filled with ACSF was placed within 20 µm of a proximal apical dendrite (50-100 µm from the soma) in layer 2/3 neurons. Episodic evoked fluorescence measurements were made over 5 min in ACSF, as well as during and following bath application of 20 µM carbachol. In the early experiments, fluorescent illumination was provided by a 100W halogen light source, and a Sutter Lambda-2 filter wheel controlled the selection of excitation filters and provided shuttering. In the later experiments, illumination was provided by LEDs (470 nm for Fluo-5F and 530 nm for ACR; Thorlabs) controlled by an analog pulse from the computer. Imaging of Fluo-5F and Lucifer Yellow used a Chroma 480/40 nm bandpass excitation filter, a Q505LP dichroic mirror, and HQ510LP long-pass emission filter. Imaging of the AlexaFluor 568 and ACR used a HQ545/30 bandpass filter for excitation, a Q570LP dichroic mirror, and a HQ610/75 nm bandpass filter. A Photometrics QuantEM 512-SC CCD camera was used to image the cells and measure fluorescence transients. Imaging of soma and dendrites were carried out at ∼93 frames/second, using 8X8 binning. Illumination was provided only during the recording periods to minimize bleaching and potential photodynamic damage. Fluorescence imaging and electrophysiological recordings were synchronized in hardware. Each voltage trace consisted of baseline period of at least 50 ms, followed optionally by local fiber stimulation or evoked APs, and continued for 2.5-3 seconds. Four stimulus conditions (EPSP alone, postsynaptic action potential stimulation alone, and combined EPSP and action potential stimuli with +10 and +50 ms intervals) were interleaved every 5-7 seconds throughout the protocol, including during and following drug delivery. As no clear calcium signals were detected under the EPSP-alone condition, those traces were used to compute a linear bleaching correction estimate that was applied to the traces for all other conditions.

A single structural image was obtained from the AlexaFluor 568 or Lucifer Yellow fluorescence before and after each run, and used to place ROIs along the visible primary apical dendrite and along the proximal regions of the first secondary branches. Changes in fluorescence, ΔF/F (= (F(t)-F_0_)/F_0,_ where F_0_ is the baseline fluorescence prior to stimulation and F(t) is the time course of the fluorescence change) were then computed, and summarized as the area under the curve of the evoked calcium-dependent fluorescent signal. Fluorescence traces for bursts of action potentials, with or without preceding EPSPs, are averages of ∼10 traces measured with the same ROI. Off-line data analysis for the imaging was carried out using Python scripts under ACQ4. Traces with spontaneous action potentials before or after the stimulus train, EPSP-evoked action potentials, or inconsistent AP production during the train, were excluded from analysis. Under our optical conditions, dendritic spines were not clearly resolved in the binned images, in which each pixel was a 2×2 µm square. Consequently, the reported measurements are likely dominated by the larger fluorescence signals from the dendritic shaft, with a smaller contribution from spine calcium signals. All of the ROIs for a given cell were scanned to determine which one showed the largest increase in the calcium signal when comparing the +50 ms EPSP-AP condition with the APs alone, as described in the results. This ROI was then used for all further analysis for a given cell.

### Chemicals and Pharmacological Agents

Carbachol, eserine, oxotremorine-M, pirenzepine, 4-DAMP, D-APV, CNQX, VU-0255035, and BAPTA were purchased from Tocris. AlexaFluor 568, Lucifer Yellow CH, and Fluo-5F were purchased from Molecular Probes and Invitrogen. Asante Calcium Red was purchased from TEFLABS. All other chemicals were purchased from Sigma-Aldrich.

### Statistical Analysis

Data are reported and plotted as means and sample SD. Statistical comparisons were made using one or two-way ANOVA (with post-hoc tests using Holm-Sidak’s multiple comparison corrections when specific subsets of observations are compared, or Tukey’s when all observations are being compared), paired or unpaired two-tailed t tests or single-sample t-tests (when comparing an effect at a within-cell basis for a single experimental group) as appropriate. All t tests used Welch’s correction and assumed unequal variances (Ruxton 2006). Comparisons of intrinsic parameters used a multivariate analysis of variance (MANOVA). Analysis of the calcium signals used a linear mixed-effects model fit by maximum likelihood, followed by multiple comparisons of means using Tukey contrasts. Statistics were computed using Prism (V6.0 and 7.0; two-way ANOVAs and one-way t-tests), scipy.stats (V0.17.1; t-tests with Welch’s correction), and R (V3.3.1; multiple ANOVA, linear mixed-effects models; (R Development Core Team 2018)). Statistics are reported with degrees of freedom, the value of the statistic if available, the p value, the number of cells (the unit of analysis) and the number of mice from which the cells were obtained. Statements of statistical significance are based on p <0.05, but exact p values are reported.

## Results

As indicated in the methods, the results reported here were obtained from two sets of experiments performed in different time frames by different investigators. Although most of the experimental conditions for the two sets of experiments were identical (mouse strain, anesthesia, slice preparation methods, bath solutions, electrode solutions, and temperature), the stimulation levels used in the two sets of experiments resulted in different amplitude EPSPs. In the first sets of experiments, baseline EPSPs (averaged across all cells in a single experimental condition, for the first 5 minutes of recording) averaged 7.1 mV (SD 1.5 mV; N = 24 conditions, range 4.6-10.5 mV). In the second set of experiments, baseline EPSPs across experimental groups were on average smaller at 3.1 mV (SD 1.1 mV; N = 12 conditions, range 0.9-4.8 mV). The baseline EPSP amplitudes were significantly different (t_29.8_ = 9.25, p=0.0001, two-tailed t-test with Welch’s correction), as were the initial EPSP slopes (t_23.2_ = 6.78, p=0.0001).

### Spike timing-dependent plasticity at layer 2/3 synapses in AC

STDP was induced by pairing EPSPs generated from stimulation in layer 2/3 with postsynaptic action potentials evoked by direct current injection through the recording electrode. Baseline EPSPs were monitored by stimulating at 0.1Hz. After 5 minutes of baseline stimulation, the STDP induction protocol was presented, after which EPSPs were monitored at 0.1 Hz for 30-40 minutes. The induction protocol consisted of an EPSP paired with 5 action potentials at 125 Hz, repeated 100 times at 1 Hz. In control experiments, only presynaptic EPSPs were elicited, only postsynaptic action potentials were elicited, or no stimulation was used during the induction protocol. The timing between the EPSP and the first (for pre-before-post pairing) or last (for post-before-pre pairing) action potential was varied to investigate timing-dependent plasticity rules. In layer 2/3 neurons, pairing of EPSPs with postsynaptic spikes resulted in bidirectional plasticity that depended on the EPSP-spike timing. Synaptic plasticity was measured by comparing the mean slope of the rising phase of the EPSP measured from 20-40 minutes after the induction protocol to the baseline slope determined from the 5 minutes prior to the pairing (S/S_0_). When spikes preceded the EPSP by 20 ms (−20 ms), no tLTP or tLTD was observed (Figure 2A, B; mean S/S_0_=0.885 (SD 0.218), t_5_=-1.184, p=0.29, N=6 cells from 4 mice, one-sample t-test). When spikes preceded the EPSP by 10 ms (−10 ms), a significant tLTD was observed (Figure 2C, D; mean S/S_0_=0.750 (SD 0.076), t_4_=-6.611 p=0.0027, N=5 cells from 4 mice). On the other hand, when the onset of EPSPs preceded spikes by 10 ms (+10 ms), tLTP was induced (Figure 2E, F; mean S/S_0_=1.457 (SD 0.369), t_8_=3.507, p=0.008, N=9 cells from 9 mice). No synaptic plasticity resulted when the interval between EPSP and spikes was +20 ms (Figure 2G, H; mean S/S_0_=0.984 (SD 0.363), t_6_=-0.096, p=0.93, N=7 cells from 7 mice). In contrast, EPSPs preceding spikes by 50 ms (+50 ms) resulted in tLTD (Figure 2I, J; mean S/S_0_=0.605 (SD 0.197), t_4_=-4006, p=0.016, N=5 cells from 4 mice). A separate group of cells was tested at +10 ms in the second set of experiments (Figure 3A, unfilled circles at +10 ms), slightly weaker stimulation that resulted in smaller EPSPs (mean 4.79 mV (SD=1.33), S/S_0_=2.67 (SD 1.364), N = 6 cells) compared to the EPSPs in the first data set (mean 5.66 mV (SD=1.91), N=9) (filled circles, Figure 3A at +10 ms). Although the second group did not show a statistically significant effect of the induction protocol (mean S/S_0_=1.716 (SD 0.855), t_5_=1.874, p=0.12, N=6 cells from 2 mice; 4 male and 2 female), the mean effect size was larger than in the first group. Furthermore, no significant difference in the EPSP slope ratios was observed between these two groups (t_9.84_=-0.64, p=0.544; unpaired t-test with Welch’s correction). Combining the two data sets still shows a significant effect at +10 ms (mean S/S_0_=1.561 (SD 0.625), t_14_=3.361, p=0.0047, N=15 cells from 11 mice). In subsequent comparisons, these two sets of measurements at +10 ms are used separately as controls for contemporaneous manipulations.

**Figure 3:**
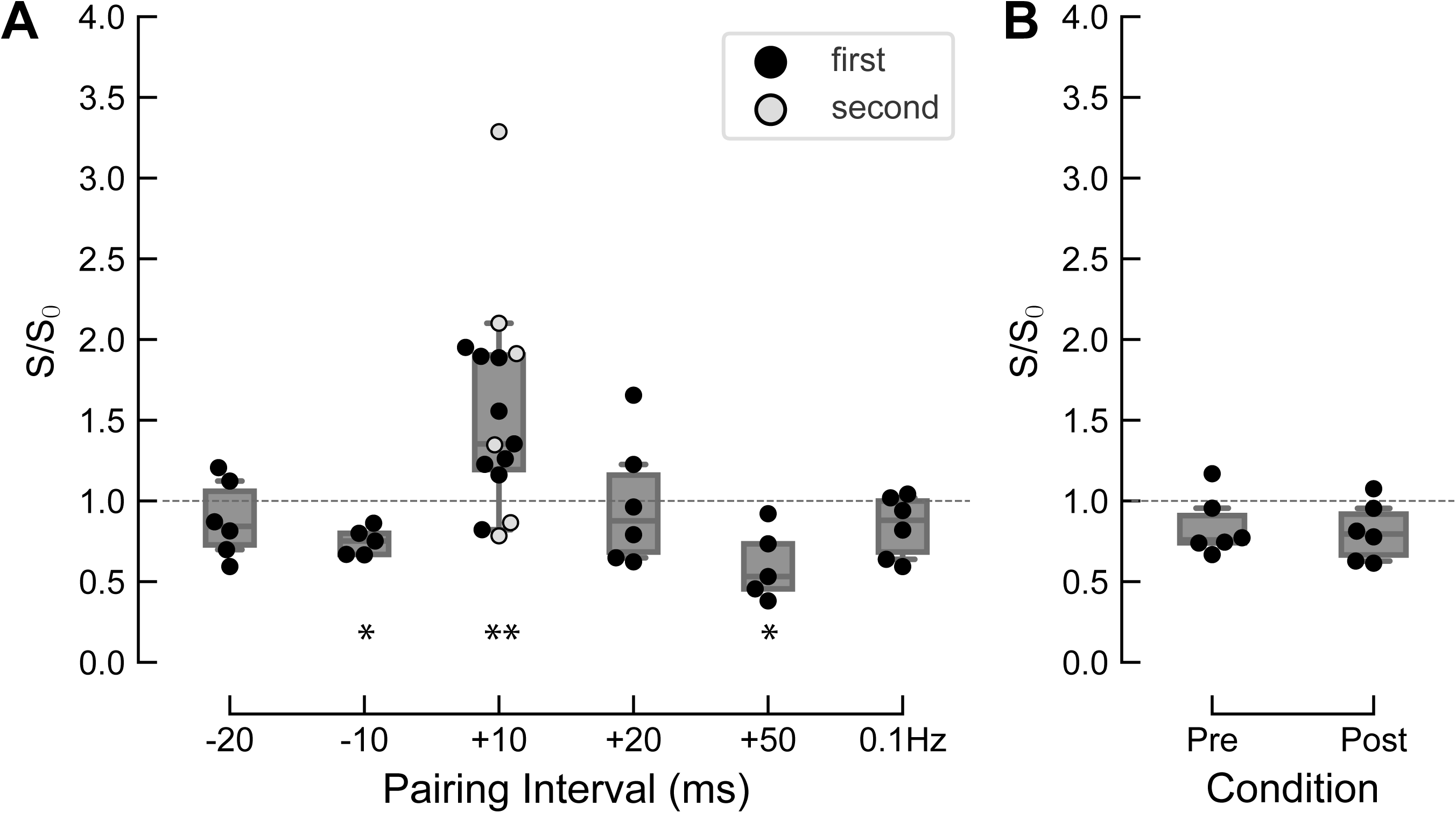
Summary of changes in synaptic strength across pairing intervals and for control conditions. A. Summary of EPSP slopes across pairing intervals, and a low-frequency pairing control (0.1 Hz). Each point is an individual cell; boxes show median and interquartile distances; whiskers show 5 and 95% confidence intervals. Black circles are data from the first set of experiments; gray circles are from a second independent set of experiments at the +10 ms interval (see text). The single asterisks indicate significant depression at −10 and +50 ms (p < 0.05) when compared to the +10 ms interval; double asterisks indicate significant (p<0.005; combined datasets) tLTP at the +10 ms interval (1-way ANOVA, followed by Tukey’s test for all pairwise combinations of intervals). B. EPSP slopes showed a slight depression with either no postsynaptic action potentials during the paring period (“Pre” only condition), or only action potentials and no EPSPs during the pairing period (“Post” only condition).

The resulting STDP curve is summarized in Figure 3A. The pairing of EPSPs and spikes had a significant interval-dependent effect (one-way ANOVA, F_5, 40_=4.780, p=0.0035). Although individual comparisons between baseline and post-STDP induction responses showed significant effects at +10, +50 and -10 ms, post-hoc comparisons between all intervals using Tukey’s multiple comparison test showed that the +10 ms interval was significantly different from both the +50 ms interval (p=0.012) and the -10 ms interval (p=0.049); all other interval pairs had p-values >0.1. Although some run-down was visible, the presynaptic stimulation-only, postsynaptic action potentials-only and 0.1Hz control (Figure 3B) were not significantly different from their baselines (0.1Hz: mean=0.837 (SD 0.182), t=−2.195, p=0.080, N=6 cells from 4 mice; pre-only: mean=0.833 (SD 0.187), t=−2.194, p=0.080, N=6 cells from 5 mice; post-only=0.810 (SD 0.184), t=−2.537, p=0.052, N=6 cells from 4 mice). The general shape of the STDP curve is similar to the curves reported at other synapses, including the presence of tLTP at short positive intervals and evidence for a weaker tLTD at short negative intervals. The presence of tLTD at +50 ms appears to be an unusual feature. The LTP at +10 ms is associative, as the induction requires both specific timing and an appropriate temporal order between pre- and postsynaptic activity.

### mAChR activation induces LTD of synaptic potentials at layer 2/3 to layer 2/3 synapses in AC

Previously, it was shown that activation of mAChRs, via electrical stimulation in layer 6 or the underlying white matter, induces a LTD of synaptic potentials in layer 3 pyramidal cells in AC (Kaur et al. 2005). To test whether mAChR activation at layer 2/3 to layer 2/3 synapses causes LTD, we bath-applied the cholinergic receptor agonist carbachol (20 µM) for 5 minutes while measuring EPSPs in layer 2/3 neurons. Carbachol induced a large transient depression of EPSPs during drug application (Figure 4A), to S/S_0_=0.528 (SD 0.088) (t_5_=-11.94, p=0.000073, N=6 cells from 2 mice, one-sample t-test compared to the normalized baseline of 1.0), measured immediately after the drug application. EPSP amplitudes eventually returned to baseline over 20 minutes. We also tested the muscarinic receptor agonist oxotremorine-M (3µM; Oxo-M). Oxo-M (Figure 4B) also produced a significant transient depression (10-25 minutes to a mean S/S_0_ of 0.597 (SD 0.127), t_4_=-6.34, p=0.0032, N=5 cells from 4 mice); when combined with the second set of experiments, the mean S/S_0_ was 0.632 (SD 0.166), t_4_=-5.864, p=0.00062, N=8 cells from 7 mice; 2 cells were from 2 male mice; the remainder had undetermined sex). Endogenous activation of AChRs with eserine, a cholinesterase inhibitor (1µM) induced a weak transient depression of EPSPs but the effect was not significant (Figure 4C, mean S/S_0_=0.812 (SD 0.188), t_5_=-2.241, p=0.075, N=6 cells from 4 mice). To further confirm that the carbachol-induced LTD measured at synapses in layer 2/3 requires activation of mAChRs rather than nicotinic acetylcholine receptors, the nonselective mAChR antagonist atropine was applied at 10 µM, a concentration that blocks all mAChR subtypes (Peralta et al. 1987), prior to and during the application of 20µM carbachol. Atropine blocked the transient depression produced by carbachol (Figure 4D, mean S/S_0_ from 10-25 minutes = 0.883 (SD 0.107), t_2_=-1.543, p=0.27, N=3 cells from 2 mice). One member of the mAChR family, the M1 receptors, are highly expressed in cortex (Hohmann et al. 1995; Roβner et al. 1993). We therefore next tested two M1 receptor antagonists, pirenzepine and specific M1 competitive antagonist VU0255035 (Sheffler et al. 2009) for their ability to block the effects of carbachol. Pirenzepine at 75nM, a concentration that blocks M1 receptors somewhat selectively (Buckley et al. 1989) did not prevent the depression produced by carbachol (Figure 4E, mean S/S_0_=0.550 (SD 0.155), t_3_=-5.04, p=0.015, N=4 cells from 2 mice). Because VU0255035 was solubilized in DMSO prior to addition to the ACSF, an additional set of control experiments with 20µM carbachol and 0.05% DMSO was performed in a separate set of cells (Figure 4F; 8 cells from 4 mice), and the results were only compared between the two groups. As shown in Figure 4G, 5 µM VU0255035 significantly reduced the depression produced by carbachol, from 0.339 (SD 0.131, N = 8 cells from 4 mice) to 0.641 (SD 0.190, N=5 cells from 2 mice, t_10.80_=-3.12, p=0.019). Comparing these manipulations (Figure 4H; excluding the VU0255035 experiments because they were done under different conditions), including the 0.1 Hz no-drug conditions (Figure 3B), for the period starting 5 minutes after the onset of drug application through 5 minutes following application (a 15-minute period) reveals a significant effect of treatment (F_5, 24_=4.59, p=0.0044). Post-tests were done to compare the effects of the drug treatments against the 0.1Hz data, and to test the effects of the antagonists against the carbachol effect. Compared to the 0.1Hz, carbachol (p=0.016) resulted in depressed EPSPs, while depression was not significant in the presence of Oxo-M (p=0.11). Atropine blocked the effect of carbachol alone (p=0.025), whereas neither eserine (p=1.0) nor 75 nM pirenzepine had a discernible effect (p=1.0 in each case).

**Figure 4:**
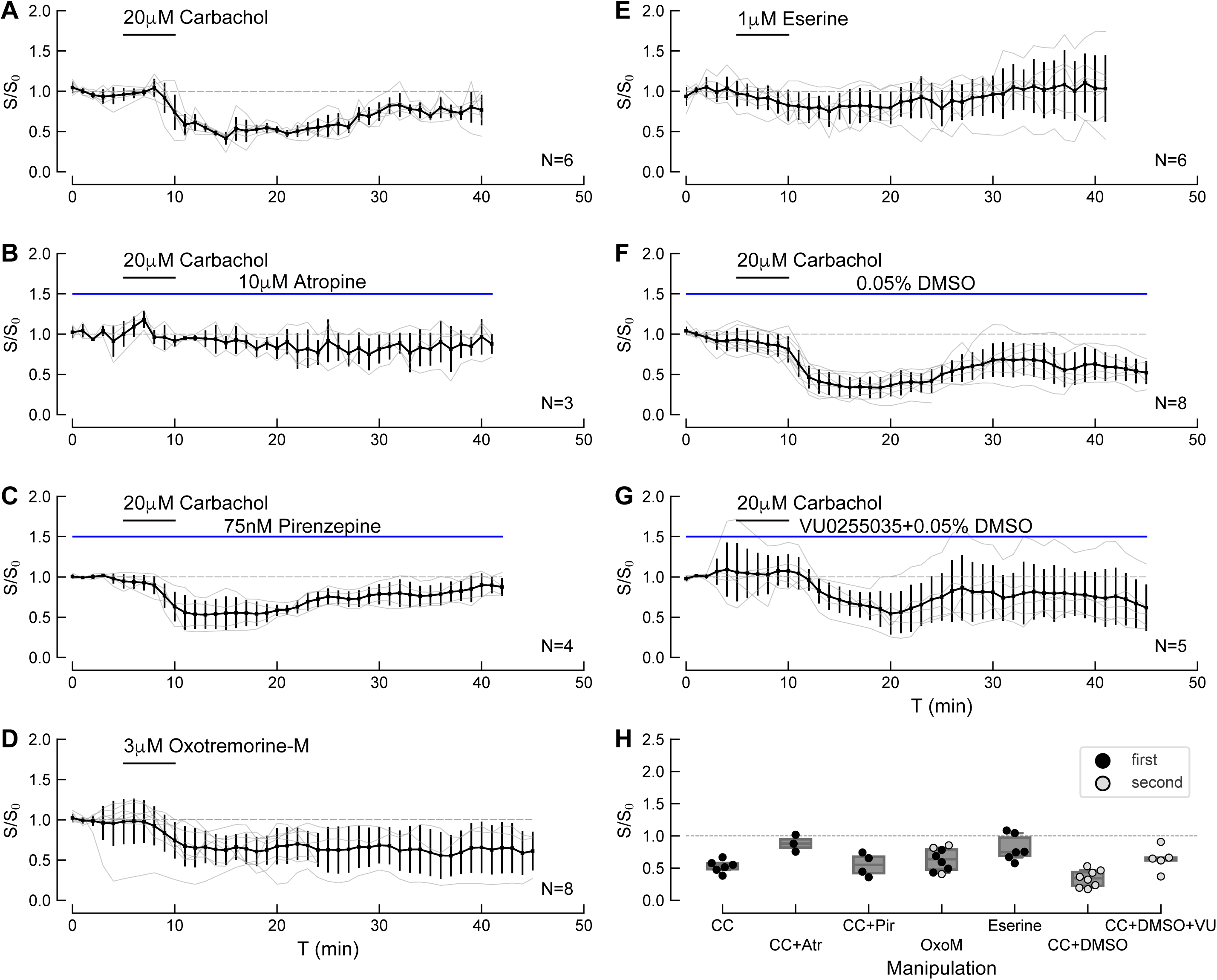
Muscarinic receptor activation depresses synaptic transmission at layer 2/3 synapses onto auditory cortical pyramidal cells. A. The cholinergic agonist carbachol (20 µM, 5 mins) elicits a transient depression during agonist application followed by a weak but continuing LTD after washout that lasts the duration of the recording. B. The carbachol-induced transient depression is prevented by 10 µM atropine, a nonselective mAChR antagonist. C. Carbachol-induced depression is not inhibited by 75 nM pirenzepine. D. The muscarinic receptor-specific agonist, Oxotremoroine-M, produces a transient depression, similar to that produced by carbachol. E. Application of the anticholinesterase eserine (1 µM, 5mins) induces a weak reversible transient depression. F. Depression of transmission with 20 µM carbachol is not affected by 0.05% DMSO. Data are from a separate set of cells than those shown in panel A. G. The M1 receptor-specific antagonist VU0255035 blunts the effects of carbachol (compare to panel F; p < 0.02; unpaired t-test). A-G. Error bars are SDs, and the time of drug application is shown by the horizontal bar in each graph. N indicates the number of cells in each data set. The time course of EPSP slope for individual cells is shown by the faint colored lines. H. Summary of slopes measured from average EPSPs for the first 10 minutes after drug application. Each point is the measurement for a single cell; light grey points with a black outline are from data in the second set of experiments. Boxes indicate the median and interquartile distances.

We next tested whether the carbachol-induced transient depression of transmission was induced pre- or postsynaptically, by examining the paired pulse ratio (PPR) of EPSP slopes. Although not definitive, a change in PPR suggests a presynaptic locus of expression, whereas no change is an indicator of a postsynaptic locus (Dobrunz and Stevens 1997). Carbachol application did not change the PPR (control: 1.16 (SD 0.23); carbachol: 1.32 (SD 0.23), t_4_=1.921, p =0.13, paired t-test, N=5 cells from 4 mice), suggesting that a postsynaptic mechanism underlies the reduction in EPSP size produced by carbachol. Taken together, these results suggest that exogenous activation of mAChRs causes synaptic depression at synapses onto layer 2/3 pyramidal cells, and that these effects occur through postsynaptic mechanisms. A portion, but likely not all, of this effect may be mediated by M1 receptors.

### Modulation of intrinsic excitability

Spike-timing dependent plasticity can also be affected by changes in intrinsic excitability. We therefore measured the effects of carbachol and Oxo-M alone on current-evoked firing and action potential shape. Figure 5A shows spiking of a layer 2/3 pyramidal cell in response to current injections under control conditions. Following 5 min incubation with carbachol (20µM), the cell fired more rapidly in response to the same currents (Figure 5B). Equation 1 was fit to the the FI curves to extract the rheobase (I_break_), the rate of firing increase with current, and the maximal firing rate. Carbachol (20µM, Figure 5C) caused a reversible increase in excitability measured as an enhancement in firing rate in response to current steps (I_rate_: control: 0.0044 (SD 0.0016) pA; carbachol: 0.0120 (SD 0.0025) pA; t_10.62_=−4.327, p=0.0019, N=10 cells from 5 mice, paired t-test with Welch’s correction). Neither the maximal firing rate F_2_ (control: 34 (SD 29) sp/s; carbachol: 17 (SD 5) sp/s; t_9.49_=2.16, p=0.059, N=10), or the spike threshold I_break_ (control: 25 (SD 18) pA; carbachol: 32 (SD 24) pA; t_16.87_=−1.407, p=0.19, N=10) were altered by carbachol. The adaptation ratio during carbachol was significantly decreased (control: 3.928 (SD 1.196); carbachol: 2.936 (SD 0.587); t_13.09_=3.278, p=0.0096, N=10), and the membrane potential depolarized by a small amount (control: -63.2 (SD 1.6) mV; carbachol: -62.2 (SD 1.2) mV; t_17.03_=−2.443, p=0.037, N=10). The mean first action potential half-width increased by ∼0.5 ms, but this effect was not significant (control: 1.31 (SD 0.31) ms; carbachol: 1.89 (SD 1.13) ms; t_10.31_=−1.652, p=0.13, N=10). The input resistance also did not change (control: 254 (SD 61) MΩ; carbachol: 249 (SD 85) MΩ; t_16.28_=0.272, p=0.79, N=10).

**Figure 5.**
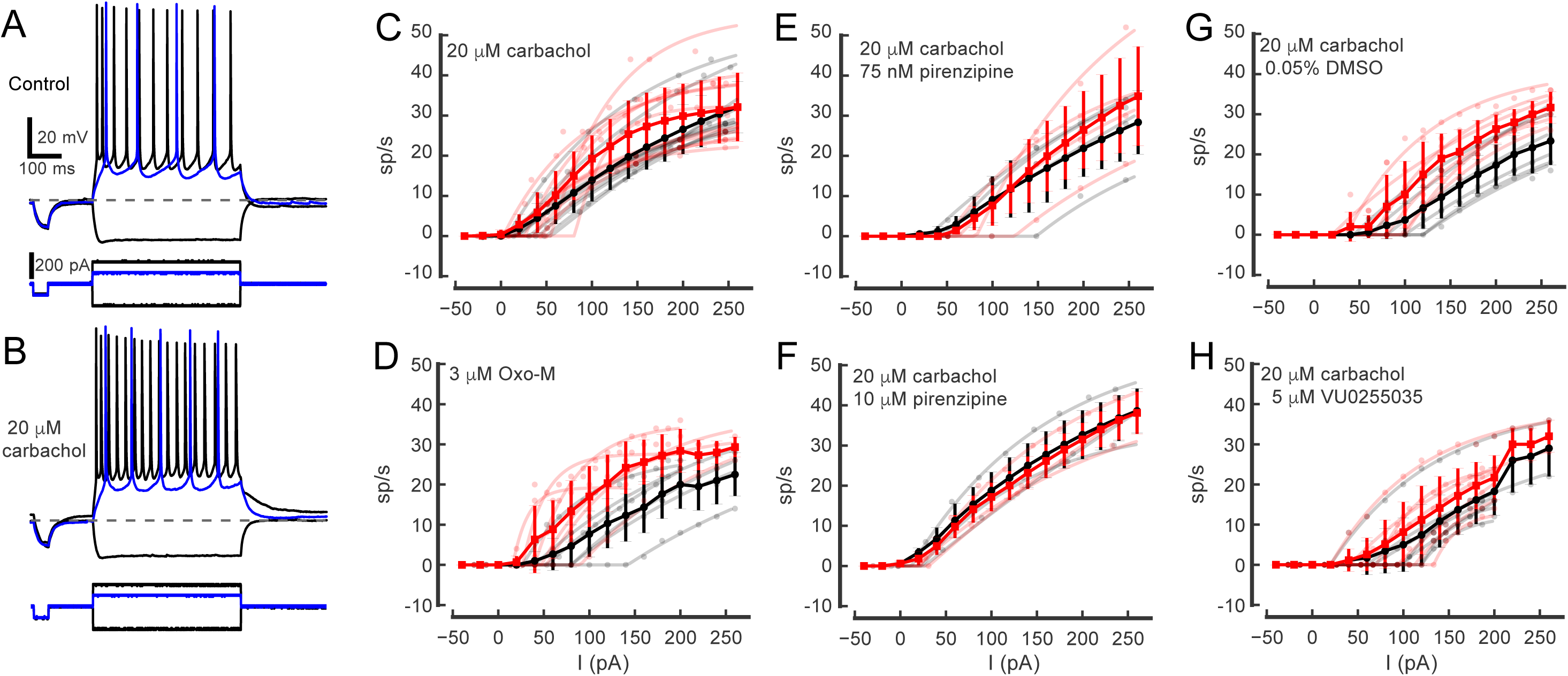
Summary of pharmacological activation and receptor block on excitability of layer 2/3 pyramidal cells. A. Responses of a cell to injections of current in control conditions with a +100 pA pulse (blue traces) and ±200 pA pulses (black traces). B. Responses of the same cell to the same current pulses in the presence of 20 µM carbachol. Carbachol increases the firing rate at both depolarizing current levels. C. Summary of firing rate with current level for control (black) and in the presence of carbachol (red). Carbachol increases the slope of the FI curve. D. Application of Oxotremorine-M (3 µM) also increases the firing rate with increasing current. E. Pirenzepine (75 nM) reduces the effect of carbachol. F. A higher concentration of pirenzepine (10 µM) completely blocks the effects of carbachol. G. Control showing increased firing in to 20 µM carbachol in the presence of DMSO. H. The M1-receptor specific antagonist VU0255035 blocks the effects of carbachol.

We also tested how Oxo-M affected the FI curves (Figure 5D). Oxo-M decreased the threshold current (I_break:_ control 110 (SD 101) pA; Oxo-M: 60 (SD 41) pA; t_10.87_=2.870, p=0.028, N=7 cells from 6 mice; 2 cells were from 2 male mice; the remainder were of undetermined sex). Oxo-M produced a small and non-significant effect on I_rate_ (control: 0.0068 (SD 0.0042) pA; Oxo-M: 0.0213 (SD 0.0184) pA; t_6.64_=−1.971, p=0.096, N=7). A similar small but non-significant effect was evident in the maximal firing rate, F_2_ (control: 20 (SD 6) sp/s; Oxo-M: 15 (SD 5) sp/s; t_9.24_=1.908, p=0.105, N=7). Interestingly, Oxo-M depolarized cells by approximately 6 mV on average (control: -67.7 (SD 3.9) mV; Oxo-M: - 61.3 (SD 4.4) mV; t_11.82_=−3.441, p=0.014, N=7) and produced an increase in action potential half width (control: 1.26 (SD 0.26) ms; Oxo-M: 1.59 (SD 0.27) ms; t_11.93_=−3.293, p=0.016, N=7). No other parameters were significantly affected (all p > 0.09). Overall, these descriptive analyses suggest that activation of muscarinic receptors increases the excitability of the cells through a combination of weak depolarization and an increased steepness of the firing rate with current.

To further understand how mAChR activation affects excitability, we explored the pharmacology of receptor activation on the firing of layer 2/3 pyramidal cells. Pirenzepine (75 nM; Figure 5E) appear to blunt the effects of carbachol, as there were no significant changes in RMP, adaptation ratio, first action potential half-width, or the fitted parameters of the FI curves, I_break_, F_2_, and I_rate_ (all p > 0.18, Welch’s t-test, N=4 cells from 2 mice). R_in_ was lower in the presence of pirenzepine however (carbachol alone: 208 MΩ (SD 65), carbachol + pirenzepine: 160 MΩ (SD 71); t_5.94_ = 4.471, p=0.021). Similarly, 10 µM pirenzepine (Figure 5F) blocked all the effects of carbachol (all p > 0.23, Welch’s t-test, N=3 cells from 3 mice). Likewise, the more specific M1 receptor antagonist VU0255035 largely prevented the shifts seen with carbachol. As this drug was solubilized in 0.05% DMSO, we performed a separate set of control experiments with carbachol in the presence of DMSO for comparison. Carbachol in DMSO produced a similar increase in excitability as under control conditions (Figure 5G). I_break_ was lower (control: 90 (SD 32) pA; carbachol+DMSO: 66 (SD 32) pA; t_9.99_=4.756, p=0.0051, all comparisons are paired t-tests, N=6 cells from 2 mice; 1 cell was from a male mouse and the remainder were of undetermined sex), and I_rate_ was higher (DMSO alone: 0.0037 (SD 0.0012) pA; carbachol+DMSO: 0.0096 (SD 0.0017) pA; t_7.73_=−7.224, p=0.00079) in carbachol, but F_2_ was not different (DMSO alone: 27 (SD 11) sp/s; carbachol+DMSO: 20 (SD 3) sp/s; t_5.62_=1.905, p=0.12). As with carbachol alone and Oxo-M, the membrane potential depolarized (DMSO alone: -67.4 (SD 4.1) mV; carbachol+DMSO: -62.7 (SD 3.2) mV; t_9.40_=−5.925, p=0.0020), and the adaptation ratio decreased (DMSO alone: 2.599 (SD 0.648); carbachol+DMSO: 1.989 (SD 0.561); t_9.80_=6.389, p=0.0014). A small but non-significant increase in the first action potential half-width was also seen (DMSO alone: 1.11 (SD 0.17) ms; carbachol+DMSO: 1.22 (SD 0.24) ms; t_8.85_=−2.400, p=0.062). In the presence of 5µM VU0255035, the changes in firing produced by carbachol were blocked (Figure 5H). None of the FI parameters were significantly different from carbachol in DMSO control when carbachol was tested in the presence of VU0255035 (I_break_: (carbachol-DMSO: 83 (SD 36) pA; carbachol-VU0255035: 75 (SD 40) pA; t_11.88_=1.561, p=0.17, all comparisons are paired t-tests, N=7 cells from 3 mice; 1 cell was from a male mouse and the remainder were undetermined); F_2_: (carbachol-DMSO: 21 (SD 14) sp/s; carbachol-VU0255035: 18 (SD 10) sp/s; t_10.50=_0.863, p=0.42), I_rate_ (carbachol-DMSO: 0.0126 (SD 0.0184) pA; carbachol-VU0255035: 0.0101 (SD 0.0114) pA; t_11.88_=−0.403, p=0.70). The adaption ratio also did not change (carbachol-DMSO: 1.927 (SD 0.791); carbachol-VU0255035: 1.993 (SD 0.726); t_11.91_=−0.579, p=0.58). However, the resting membrane potential still showed a modest depolarization (carbachol-DMSO: -64.1 (SD 4.6) mV; carbachol-VU0255035: -61.6 (SD 3.2) mV; t_10.60_=−2.834, p=0.030, N=7 cells), and the action potential half width was significantly wider (carbachol-DMSO: 1.22 (SD 0.21) ms; carbachol-VU0255035: 1.42 (SD 0.32) ms; t_10.34_=−3.034, p=0.023). Using MANOVA to compare all excitability measures with a carbachol challenge in the presence of VU0255035 against the carbachol challenge in DMSO however revealed no difference between the two groups (F_7,5_, p=0.083). Individual post-hoc comparisons only showed a significant difference in the adaptation ratio (F_1,11=_14.48, p=0.0029). Taken together these data suggest that activating the muscarinic receptors with either carbachol or Oxo-M increases overall excitability of the cells and in increases the steepness of the FI curve. Blocking the mAChRs seems to partially prevent these changes, but the effects are modest and not consistent between 10 µM pirenzepine and VU0255035, suggesting that the two compounds likely have different profiles with respect to antagonism of carbachol at different receptor subtypes that are differentially coupled to the ion channels regulating excitability.

### mAChR activation regulates tLTP

We next tested the hypothesis that cholinergic neuromodulation by mAChRs can change STDP timing rules by modulating the strength of tLTP. We focused on the +10 ms pre-before-post intervals because the largest changes in EPSPs were seen with this timing, and we were more confident that the effects of any manipulations would not be caused by rundown of synaptic transmission. Activating mAChRs with carbachol reduced tLTP at +10ms, (Figure 6A; control: 1.457 (SD 0.391), N = 9 cells from 9 mice; same control group as above, carbachol: 0.924 (SD 0.440), N=6 cells from 6 mice, t_14.60_=2.40, p=0.037, Welch’s t-test for unpaired samples). The second set of experiments with weaker test EPSPs showed a similar suppression of the tLTP, but was not significant (mean S/S_0_=1.716 (SD 0.936), N = 6 cells from 2 mice, compared 1.309 (SD 0.384), N=8 cells from 5 mice, t_6.24_=-1.001, p=0.352, Welch’s t-test for unpaired samples). Combined, these two sets of experiments showed an potential effect of carbachol (control: S/S_0_ =1.561 (SD 0.646), N = 15 cells from 11 mice; in carbachol: S/S_0_ = 1.144 (SD 0.430, N = 14 cells from 11 mice), t_25.11_=2.05, p=0.052). To more specifically activate mAChRs, we also tested Oxo-M. Application of Oxo-M (3 µM) during induction clearly prevented tLTP (Figure 6B, mean S/S_0_=0.917 (SD 0.445), N=7 cells from 4 mice, t_15.07_=2.54, p=0.026, compared to control +10ms). To explore whether intrinsic cortical acetylcholine could modulate STDP in auditory cortical neurons, we also examined the effects of the anticholinesterase, eserine. Application of eserine during pre-before-post pairing also prevented tLTP induction (Figure 6C, mean S/S_0_=1.005 (SD 0.283), N=6 cells from 3 mice, t_15.77_=2.59, p=0.022) compared to control at +10ms), indicating that endogenous acetylcholine can prevent tLTP at excitatory synapses onto layer 2/3 pyramidal neurons. As the postsynaptic action potentials were elicited by trains of brief current pulses (and therefore controlled), the effects of mAChRs on the firing rate and cell excitability (Figure 5) were not responsible for these differences, although increases in action potential width could contribute to changes in STDP. Taken together these experiments indicate that tLTP can be modulated by activation of mAChRs through carbachol, Oxo-M or endogenours ACh.

**Figure 6:**
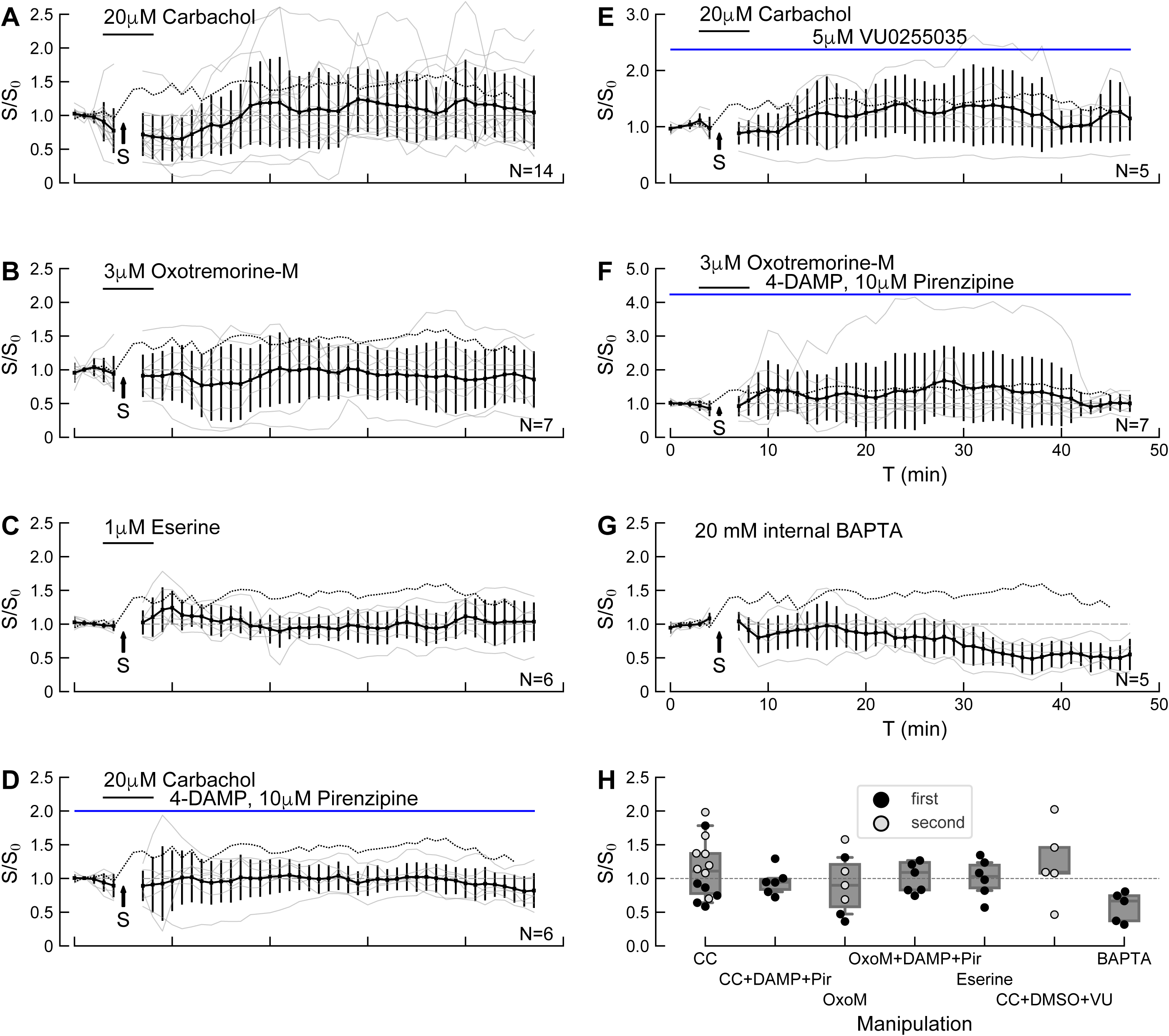
mAChR activation modulates STDP. A. Summary of effects of carbachol (20µM) on pre-before-post pairing STDP at +10ms, as in Figure 2A. Carbachol (20 µM) reduces tLTP. Solid line shows mean; error bars are 1 SD. The dashed line shows tLTP at +10 ms in control conditions (from Figure 2A). Gray lines show individual cells. B. The mAChR specific agonist Oxo-M also reduces tLTP. C. Eserine (1µM) likewise reduces tLTP after at +10ms. D. 4-DAMP and pirenzepine appear to not reverse carbachol’s suppression of tLTP, but the effect is not significant. E. The M1-specific antagonist VU025035 does not prevent carbachol’s effect on tLTP. F. Blocking M1 and M3 receptors with 4-DAMP and pirenzepine appears to prevent the reduction in tLTP produced by Oxo-M, but the effect is not significant. G. Chelating calcium with BAPTA prevents tLTP and induces a long-term LTD. H. Summary of EPSP slope changes for each manipulation. Boxes show mean, 25-75% and 5-95% (whiskers). Individual cells are coded according to when the data was collected (first set of experiments; solid black; second set, light gray with black outline). S/S0 is the ratio of the EPSP slope relative to the baseline slope. “S” with arrow indicates the time that the pairing protocol was applied.

In the dorsal cochlear nucleus, muscarinic receptor activation with Oxo-M converted postsynaptic tLTP to presynaptic tLTD by acting on M1/M3 receptors (Zhao and Tzounopoulos 2011). Becasue M1 receptors appear to at least partially contribute to the synaptic depression produced by carbachol (Figure 4) and the excitability of cells (Figure 5), it is possible that they can also influence other signaling pathways necessary for the induction of tLTP. We therefore tested the hypothesis that the cholinergic effects on tLTP induction in auditory cortex depended on the activation of M1 and/or M3 receptors. The suppression of tLTP at +10 ms by carbachol was not significantly affected by the simultaneous presence of the M1 antagonist, pirenzepine (10µM, a high concentration that also should also block M4 receptors), together with the M3 antagonist, 4-DAMP (1µM) (Figure 6D, mean with carbachol and antagonists: 0.951 (SD 0.197), N=6 cells from 4 mice, t_6.93_=-0.138, p=0.89, compared to carbachol alone in Figure 6A). Likewise, comparing tLTP in 20µM carbachol (including a subset of cells tested with carbachol in 0.05%DMSO, which were not different than carbachol alone) against tLTP in the same solution with VU0255035 showed that block of the M1 receptors did not restore tLTP (Figure 6E, carbachol +DMSO group: 1.309 (SD 0.384), N = 8 cells from 5 mice, carbachol+DMSO+VU0255035: 1.224 (SD 0.573), N=5 cells from 3 mice, t_10.65_=0.293, p=0.78). tLTP was partially restored by 10 µM pirenzepine and 1 µM 4-DAMP during the application of Oxo-M (the average time course of EPSP slopes after pairing is very similar to that in the absence of any drugs), but the results were quite variable and the net effect was not significant compared to OxoM’s reduction of tLTP at +10ms (Figure 6F, tLTP in presence of pirenzepine and 4-DAMP: 1.295 (SD 0.820), N=7 cells from 5 mice, t_9.25_=-1.07, p=0.31). Thus, although activation of mAChRs suppresses tLTP (Figure 6A-C), it appears that this effect is not the result of M1 receptor activation, and M3 and M4 receptors likewise are not required.

Finally, we tested whether the tLTP depended on intracellular calcium changes. When exogenous calcium chelators such as BAPTA are present in the intracellular solution, incoming calcium ions are rapidly buffered and free calcium concentration changes are strongly reduced (Tsien 1980). In the presence of intracellular BAPTA (10 mM), pre-before-post pairing at +10 ms failed to induce tLTP, and instead led to a clear LTD (Figure 6G, 0.581 (SD 0.222), N=5 cells from 3 mice, t_14.95_=5.34, p=0.00018, compared to +10 ms control). Thus, an increase in postsynaptic calcium during STDP induction appears necessary for the induction of tLTP.

We also performed additional exploratory experiments testing the effects of carbachol during EPSP-spike pairing at +50 ms and −10 ms. On average post-before-pre tLTD (−10 ms) was not significantly affected by carbachol (Figure 7A; mean 1.091 (SD 0.589), N=7 cells from 6 mice, t_4.23_=-1.51, p=0.178). Eserine (1 µM) also did not have an effect on tLTD induction at −10 ms (Figure 7B; 0.793 (SD 0.279), t_4.73_=−0.334, p=0.75, N=5 cells from 3 mice). Internal BAPTA likewise had no effect on the tLTD at −10 ms (Figure 7C, 0.665 (SD 0.196), N=4 cells from 3 mice, t_5.16_=0.81, p=0.466). Finally, pre-before-post tLTD at +50 ms was not significantly affected by carbachol (Figure 7D). Although the mean value changed from LTD (0.605) to LTP (1.225), there was a large variance between cells (SD 1.259), N=7 cells from 6 mice, t_4.34_=-1.28, p=0.245). In combination with the effects seen above at the +10 ms interval, these exploratory experiments suggest that mAChRs may modulate tLTP/tLTD differently depending on the timing intervals.

**Figure 7.**
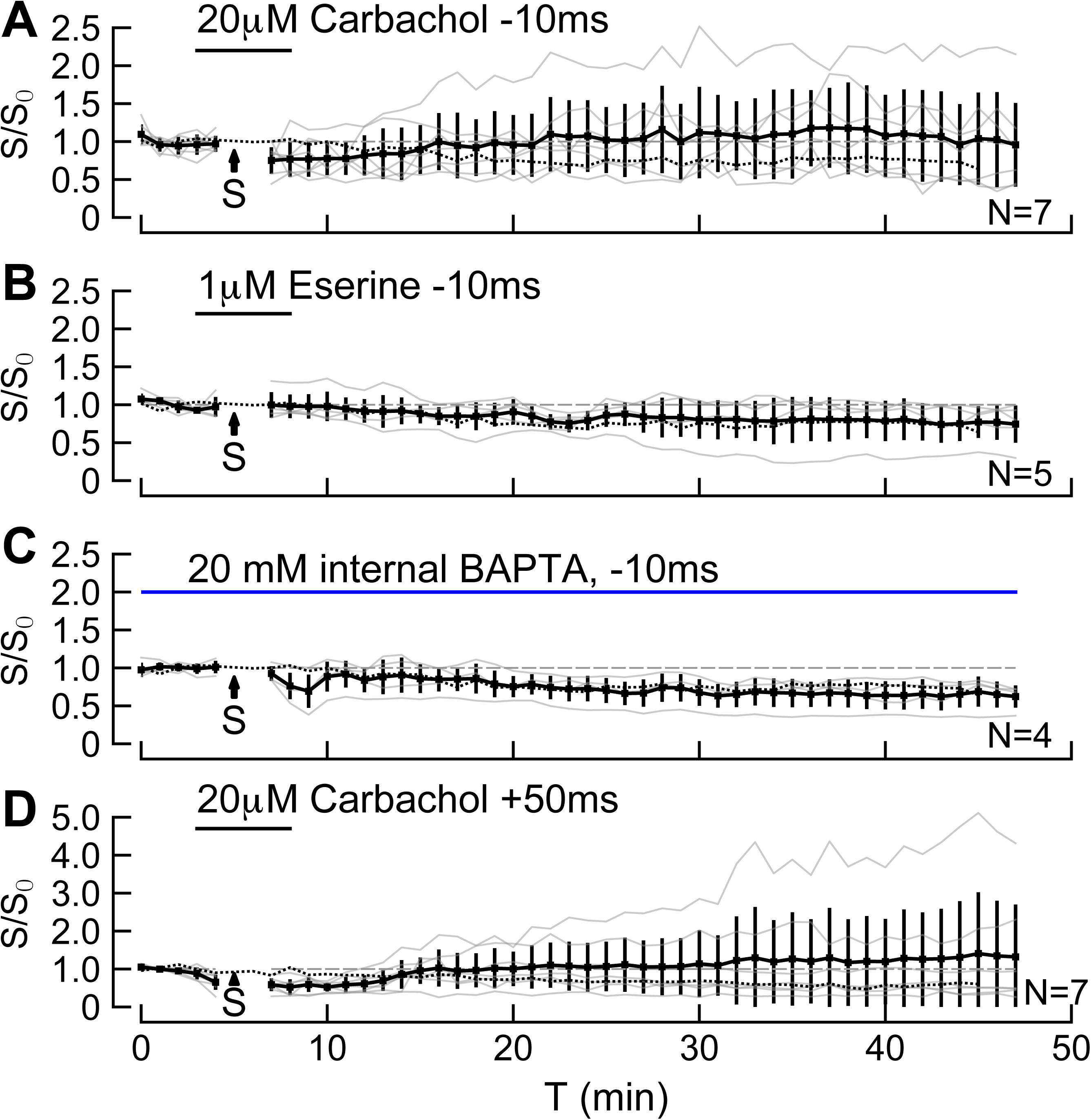
Summary of exploratory experiments at −10 and +50 ms. Data presentation format is the same as Figure 6. A. Carbachol appears to reduce tLTD at −10 ms intervals, but the effect is not significant. B. Eserine has no effect on tLTD produced at −10 ms intervals. C. 10 mM internal BAPTA has no effect on the tLTD produced at −10 ms intervals. D. Carbachol appears to prevent pre-before-post tLTP at +50 ms intervals. Dashed r lines in each panel indicate the mean of the corresponding timing STDP data from Figure 2. S/S_0_ is the ratio of the EPSP slope relative to the baseline slope. “S” with arrow indicates the time that the pairing protocol was applied.

### Activation of mAChRs reduces NMDA current

One mechanism that could account for the reduction in tLTP with mAChR (Figure 6A-C) activation is that the synaptically-evoked calcium influx through NMDA receptors is decreased and therefore does not reach the threshold required to induce tLTP. To test whether mAChR activation blocked tLTP by directly or indirectly acting on NMDA receptors, we recorded pharmacologically isolated NMDA receptor mediated EPSCs in voltage clamp with and without mAChR activation (Figure 8A). We found a reduction in the evoked NMDA receptor currents in the presence of 20µM carbachol (Figure 8B control: 263 pA (SD 123), carbachol: 148 pA (SD 104.0), t_12_=4.460, p=0.0008, N=13 cells from 6 mice, paired t-test).

**Figure 8:**
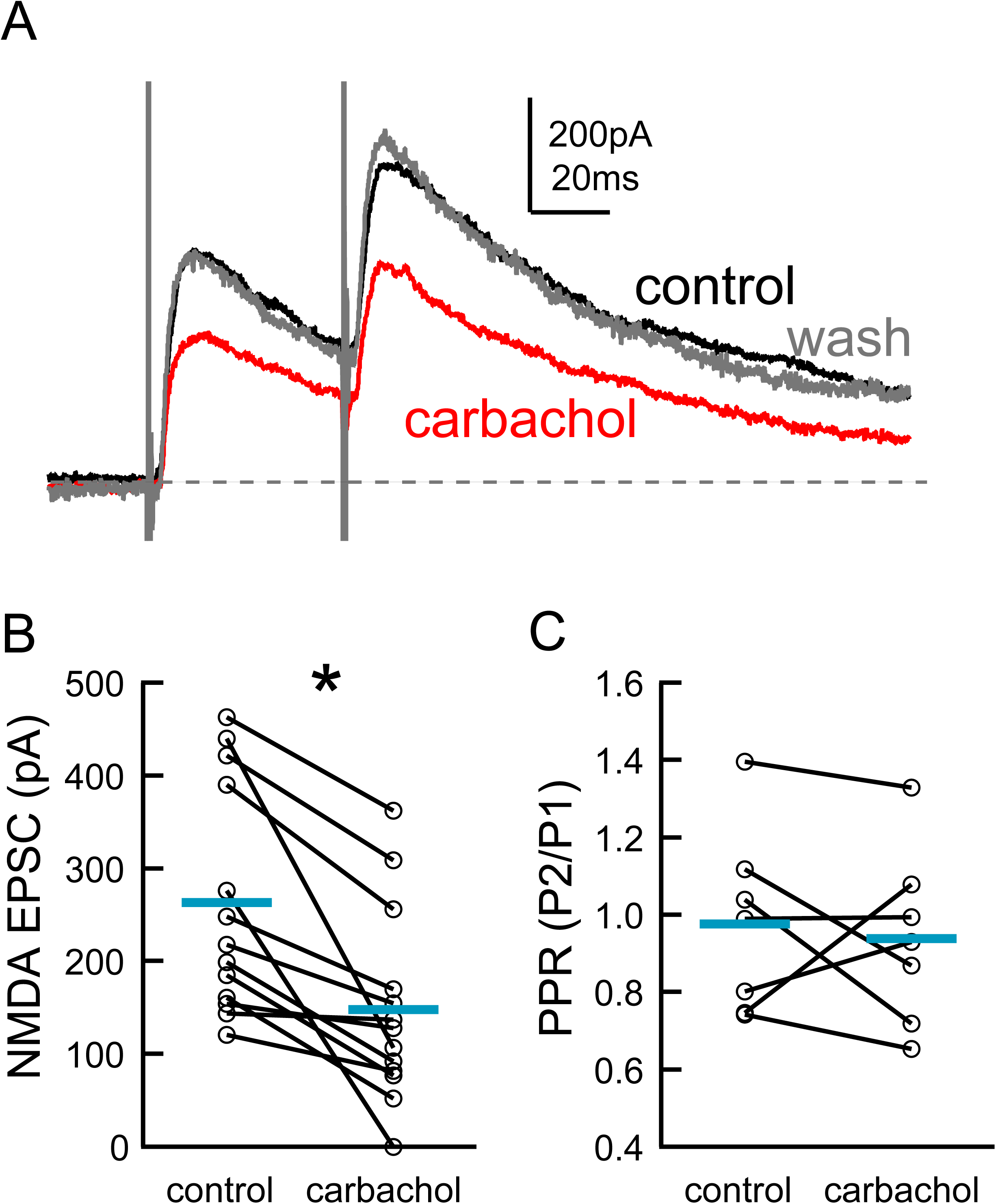
Activation of mAChRs reduces synaptically evoked NMDA currents. A. NMDA mediated synaptic currents were isolated by bathing the slice with CNQX (10µM) and picrotoxin (50µM), and cells were voltage clamped at +40mV. Carbachol (20µM, 5mins) reversibly reduced the amplitude of the isolated NMDA current. Black trace is control, red trace is carbachol, grey is after carbachol washout. B. Summary of carbachol reduction of NMDA currents (*, p<0.05, paired-t-test, N=13 cells). C. The paired pulse ratio (PPR) of the NMDA receptor currents measured at a 50 ms interval was unaffected by carbachol (p > 0.05, paired t-test, N=7 cells), suggesting a postsynaptic site of action. Horizontal blue bars in B and C show the means for each condition.

The current was partially restored after the carbachol was removed from the perfusate by replacement with normal ACSF. The current was mediated by NMDA receptors because it was nearly completely blocked following the application of the NMDA receptor antagonist D-aminophosphonovaleric acid (D-APV, 50 μM, remaining current 15.8 (SD 13.9) %, N=3; data not shown). In a subset of cells, the EPSC paired-pulse ratio was also measured to test for a potential presynaptic effect of carbachol. The paired-pulse ratio of the isolated NMDA current was not altered by carbachol application (Figure 8C, control: 0.95 (SD 0.24), carbachol: 0.94 (SD 0.23), t_6_=0.3135, p=0.76, N=7 cells from 3 mice, paired t-test) consistent with the lack of effect of carbachol on the paired-pulse ratios of EPSPs described above. Although we cannot exclude that carbachol acts presynaptically on the basis of this experiment, these results suggest that a decrease in postsynaptic current through NMDA receptors following mAChR activation could be at least partially responsible for the decrease in tLTP.

### Dendritic Calcium Signaling Is Reduced by mAChR activation

Postsynaptic calcium transients provide an associative link between synapse activation, postsynaptic cell firing, and synaptic plasticity (Koester and Sakmann 1998; Malenka et al. 1988). By definition, back propagating action potentials are essential for the induction of STDP. Because carbachol reduced the current through NMDA receptors, it is possible that it also could reduce subsequent calcium influx in dendrites of AC pyramidal cells. To investigate this, we examined postsynaptic calcium transients in the apical dendritic shafts and synapses of layer 2/3 pyramidal neurons during timed pre- and postsynaptic (action potential) activity. Figure 9A, B show the recording and stimulating configuration.

**Figure 9.**
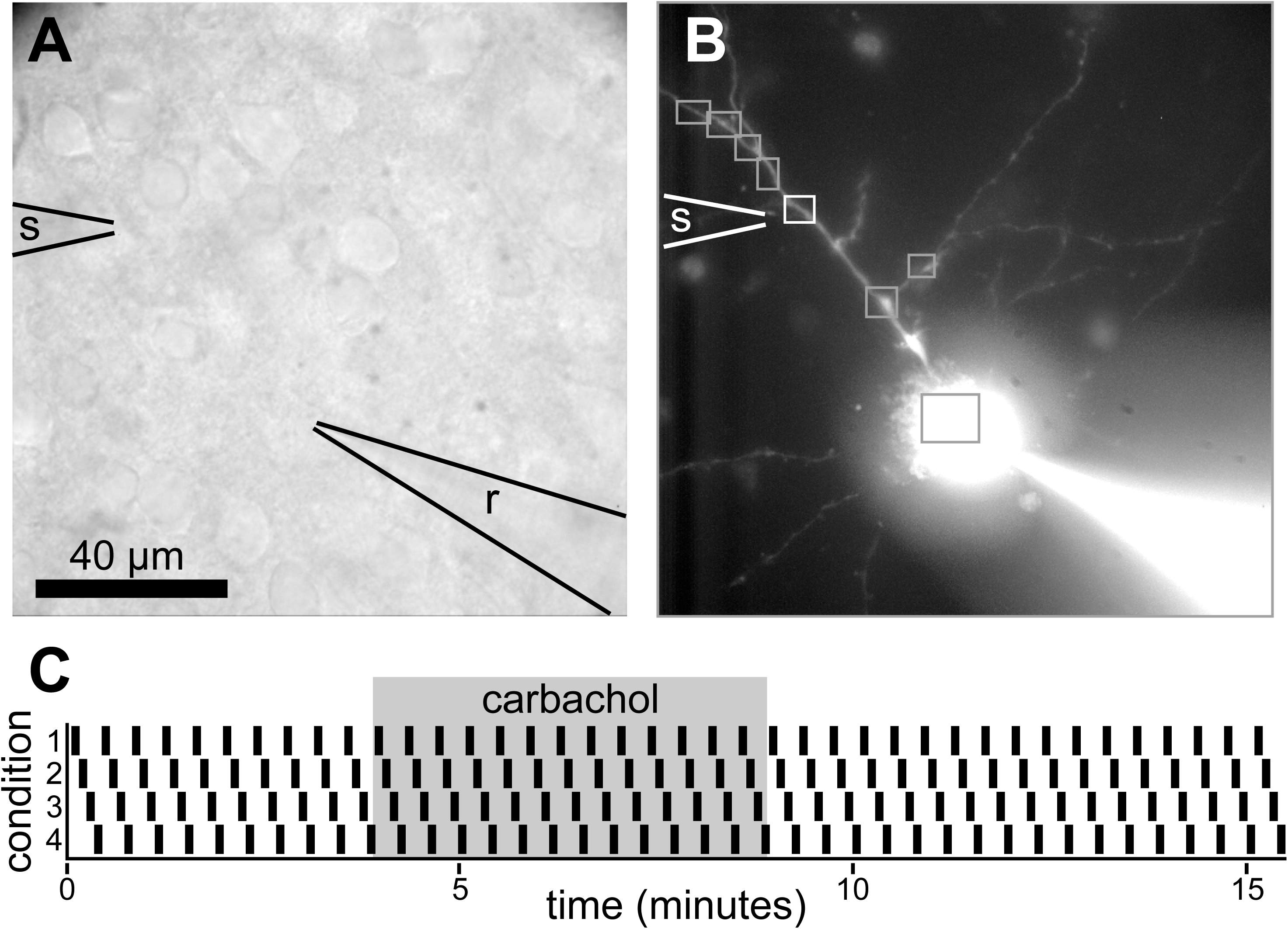
Stimulating and recording arrangement and protocols for calcium imaging experiments. A. Bright-field image showing cells in layer 2/3 and the location of the stimulating pipette and recording electrode. B. Fluorescent image of the field in A, showing a dye-filled pyramidal cell and the position of the stimulating pipette near the proximal dendrite. The white and gray boxes outline the ROIs analyzed (in an 8×8 binned image), and the white box indicates the ROI with the largest response to combined extracellular stimulation and evoked action potentials. In this case, this largest response occurred from the ROI closest to the stimulating electrode. C. Stimulation protocol pattern used in the imaging experiments. Four conditions (1: presynaptic stimulation alone, 2: presynaptic stimulation with postsynaptic action potentials at +50 ms, 3: presynaptic stimulation with postsynaptic action potentials elicited at +10 ms, and 4: postsynaptic action potentials alone) were interleaved and repeated over a 15-minute period. 20 µM carbachol was bath-applied to the slice starting at 4 minutes, and ending at 9 minutes.

Presynaptic stimulation was provided by an extracellular pipette located 50−100 µm from the soma along an apical dendrite. A series of regions along the dendrite and soma were identified to measure the calcium signals (Figure 9B) in response to a set of interleaved stimulus conditions (Figure 9C). Two sets of experiments were performed. In one set of experiments, pyramidal neurons were filled through the recording patch pipette with a structural indicator, AlexaFluor 568, and the calcium indicator Fluo-5F. The AlexaFluor 568 image was visualized and used to select a region on the apical dendrite for placement of the extracellular stimulating electrode. A burst of postsynaptic action potentials (APs) was preceded by extracellular stimulation by 10 ms, as was used for the induction of STDP. Calcium signals were analyzed in regions of interest (ROIs) placed over the primary apical dendrites and the first secondary branches. Somatically-evoked action potentials induced calcium changes throughout the visible regions of the dendritic tree. A single ROI was selected for analysis in each cell. This ROI was initially chosen as the region closest to extracellular stimulation (∼10-20 µm from electrode). All ROI’s were then tested to identify where the calcium signal in the dendritic tree was larger when EPSPs were paired with APs at the 50 ms interval than for APs alone, using a ratio measurement, with the requirement that the peak calcium signal for the AP alone be larger than 2SD of the baseline fluorescence signal prior to stimulation. In 5 of the 21 cells, this ROI was the one closest to the stimulating electrode. However, in the remainder of the cells, this ROI was elsewhere on the dendritic tree (an adjacent ROI in 7 cells, and more distant in the remainder), consistent with activation of fibers that might run vertically with layer 2/3 before contacting the target cell. All subsequent comparisons used these selected ROIs. Note that although the ROI was selected on the basis of the difference with +50 ms EPSP-AP intervals (Figure 10, A1-A3), comparisons at +10 ms and with the AP-alone condition using that ROI were based on different traces in the same cell. Although the +50 ms interval data are shown, they are not used further to avoid circularity in the analysis.

**Figure 10:**
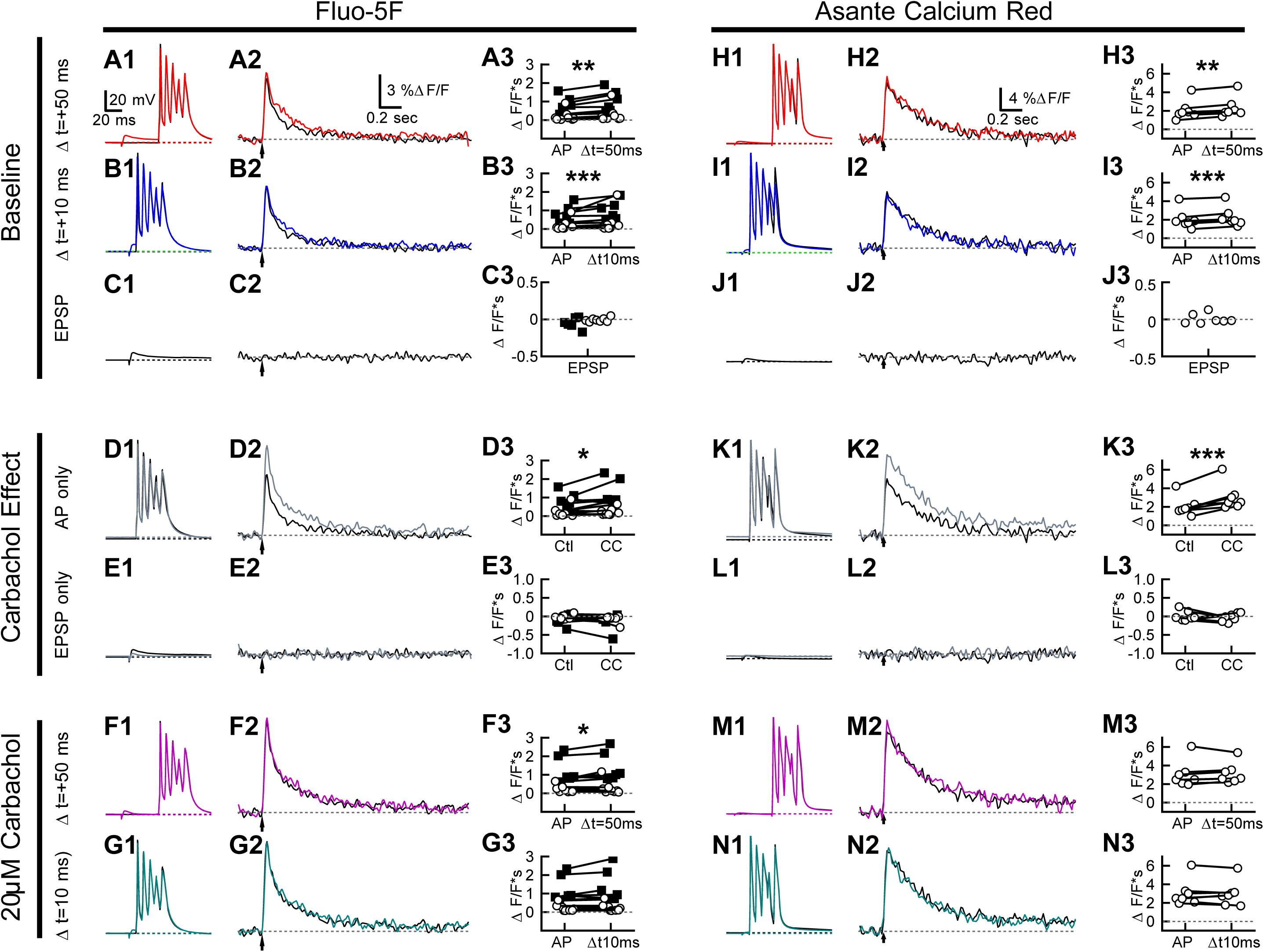
mAChR activation increases action potential evoked calcium influx in layer 2/3 pyramidal neuron dendrites. A-E. Experiments performed using the low-affinity indicator Fluo5F (N=14 cells). A1. Spike trains evoked by somatic current pulses were used to depolarize cells 50 ms after a presynaptic EPSP. Traces are average voltages during the control period (0-5 minutes in Figure 9). Black trace: no EPSP, red trace, with EPSP. A2. Calcium transients from a dendritic region of interest recorded simultaneously with the traces in A1. In the selected ROIs (see text), the pairing of the EPSP and AP produced a larger calcium transient than associated with the AP alone. A3. Summary of the integrated fluorescence signal compared between the two conditions. The fluorescence signal from the 50 ms EPSP-AP interval data was used to select the ROI that showed the largest increase with the EPSP. Each cell is consistently shown in a different color in A3-G3. See text for statistical analysis. B1-B3. Similar to A1-A3, except with a +10 ms EPSP-AP interval (blue trace with EPSP). The integrated responses in B3 for the +10-ms interval were from the same ROI’s selected in A3 for the 50-ms interval. C1-C3. EPSPs alone do not lead to detectable calcium-dependent fluorescence transients. D1-D3. APs alone produce a calcium signal that is increased in the presence of carbachol. E1-E3. No calcium signals were detected when only EPSPs were generated in the presence of carbachol. In D and E, the black traces show responses in control conditions and the grey traces show responses in the presence of carbachol. F1-F3. In the presence of carbachol, pairing EPSPs 50 ms (magenta trace) prior to APs leads to a small but significant increase in the calcium signal. G1-G3: There was no difference in the calcium signal with +10 ms pairing (dark green trace is with EPSP). All data shown in A1, A2, B1, B2, C1, C2, D1, D2, E1, E2, F1 and F2 are from one cell. H-N. Recordings from a different cell using Asante Calcium Red as the calcium indicator (N=7 cells), with the same conditions as shown in panels H-N, show a similar pattern of results. The calibration bar in A1 applies to B1-G1 and H1-N1. The calibration bar in A2 applies to B2-G2. The calibration bar in H2 applies to I2-N2. Dashed lines in A1-G1 and H1-N1 indicate the resting potential. Dashed lines in A2-N2 indicate resting fluorescence signals. Asterisks indicate statistically significant differences: *=p<0.05; **=p<0.01; ***=p<0.002 (see text for details).

We observed an increase in calcium with EPSPs and APs compared to APs alone in Fluo5F (Figure 10, B1-B3; linear mixed effects model with the integrated calcium signal and AP configuration, maximum-likelihood AIC=15.3, log likelihood=−1.63; post-tests comparing AP alone to AP+10ms EPSP: p=0.0016; AP+EPSP at +50ms: p=0.0047; N=14 cells from 7 mice; post-tests are simultaneous tests for general linear hypotheses with multiple comparisons of means using Tukey contrasts). There was no difference between the different EPSP timings (AP+EPSP at +10ms vs. AP+EPSP at +50ms: p=0.95). In response to synaptic stimulation that produced EPSPs at the soma of 2-10 mV, we were unable to resolve any changes in indicator fluorescence (Figure 10 C1-C3). A similar result was seen with ACR as the calcium indicator in a separate set of experiments (Figure 10, H-J). With ACR, cells showed an increase in the calcium signal when APs were paired with the EPSPs (AIC=21.8; log likelihood=−4.89; post-tests AP vs AP+EPSP at +50ms; p=0.0065, Figure 10, H1-H3; AP vs AP+EPSP at +10ms, p=0.0017, Figure 10 11-I3; N=7 cells from 4 mice), but no differences with EPSP timing (AP+EPSP at +10ms vs. AP+EPSP at +50ms, p=0.92). Again, there was no visible fluorescence transient with EPSPs alone (Figure 10, J1-J3).

Next, after 5 minutes of baseline measurements, carbachol (20 µM) was bath applied for 5 min. Carbachol did not induce changes in the resting indicator fluorescence. However, APs induced a much larger calcium influx in the presence of carbachol, when measured with either Fluo5F (Figure 10, D1-D3, control: 0.46 (SD 0.48, range=0.03-1.58), carbachol: 0.67 (SD 0.71, range=0.08-2.33), t_13_=−2.74, p=0.033, N=14, paired t-test) or with ACR (Figure 10, K1-K3, control: 1.47 (SD 0.72, range=0.09-2.27); carbachol: 2.38 (SD 0.74, range=1.39-3.28), t_6_=−5.17, p=0.002, N=7). The increase in AP-mediated calcium influx was blocked by atropine, implicating mAChRs (data not shown). No calcium signal was detected with EPSPs alone in the presence of carbachol with either indicator (Figure 10E1-E3 and L1-L3). Pairing EPSPs and APs produced a larger calcium influx in carbachol than without carbachol (Fluo5F: AIC=16.0, log likelihood=−2.01). In individual comparisons, the effect of EPSP+AP pairing in the presence of carbachol, compared to APs alone, was significant at the +50 ms interval (Figure 10, F1-F3, p=0.040), but not at the +10 ms interval (Figure 10, G1-G3, p=0.11) in Fluo5F. Furthermore, the calcium influx with AP+EPSP pairing was not significantly different from the AP-alone condition in carbachol when measured with ACR (Figure 10, M1-M3, p=0.41 at +50 ms; Figure 10, N1-N3, p=0.79 at +10 ms).

A linear mixed model was constructed to examine the effects of carbachol on the enhancement of the calcium signal from APs alone by EPSPs at the +10 ms interval, with a fixed effect of drug presence, and by-cell random slopes for the effects of carbachol (ROI’s for analysis of the +10 ms interval were selected using the +50 ms interval data and thus the +50 ms data were not analyzed). The two indicator dyes were treated in separate analyses. For Fluo-5F, this revealed a significant effect of carbachol, which reduced the facilitation of the calcium signal by EPSPs (F_(1, 14)_ = 9.66, p=0.0077). A similar result was observed when using ACR as the indicator (F_(1,14)_ = 5.39, p = 0.036). Taken together, these experiments indicate that, in the absence of carbachol, there is a synergistic interaction between APs and EPSPs at the 10 ms interval that results in increased dendritic calcium compared to APs alone (Figure 10, B3, I3), but this effect appears to be significantly reduced in the presence of carbachol (Figure 10, G3, N3).

## Discussion

We found that synapses activated by electrical stimulation in layer 2/3 onto layer 2/3 cells in mouse AC exhibit STDP. Although tLTP and tLTD were observed in the expected short positive (+10 ms) and negative (−10 ms) intervals, respectively, an additional tLTD was apparent at longer positive (+50 ms) intervals. We also found that mAChRs modulate tLTP and tLTD in a manner that is dependent on the EPSP-spike timing. Pharmacological activation of mAChRs when using +10 ms pairing intervals prevented tLTP induction, and could reduce tLTD at −10 and +50 ms pairing intervals, potentially leading to tLTP. tLTP at +10 ms intervals appeared to depend on intracellular calcium signaling. mAChR also activation reduced the NMDA receptor current at excitatory synapses onto layer 2/3 cells. Pairing APs and EPSPs resulted in increased dendritic calcium even when no calcium signal could be detected with EPSPs alone. This apparent supralinear calcium signal generated by pairing at +10 ms intervals was significantly decreased with mAChR activation.

### Synaptic STDP rules in AC

The magnitude and temporal structure of STDP varies with brain area, cell and synapse type (Abbott and Nelson 2000; Larsen et al. 2010). In rat AC slices, tLTP was previously observed at +10 ms intervals and tLTD at -40 ms at layer 2/3 to layer 2/3 synapses (Karmarkar et al. 2002). However, the STDP window was not further examined in that study. Other primary sensory cortical areas (V1 and S1) also exhibit before-post tLTP and post-before-pre tLTD at 10 ms intervals at layer 2/3 to layer 2/3 synapses (Froemke et al. 2006; Nevian and Sakmann 2006; Zilberter et al. 2009). A similar timing rule has been shown for synapses onto layer 5 cells in AC (D’amour and Froemke 2015). In vivo, “stimulus-timing” plasticity has also been reported in AC, by repetitively pairing acoustic stimuli (Dahmen et al. 2008) or pairing stimulation of the spinal trigeminal nucleus with acoustic stimuli (Basura et al. 2015). Within the timing range that has been examined in vivo, −30 to +30 ms, these paradigms result in tLTP and tLTD that is similar to what we report here.

In most cortical areas, the magnitude of tLTP falls off approximately exponentially with the difference between pre- and postsynaptic spike times. However, we also observed a pre-before-post tLTD at +50 ms, and on average, no tLTP or tLTD at +20 ms intervals, suggesting that the STDP curve is triphasic. Computational models have predicted triphasic STDP curves that exhibit tLTD at longer positive pre-before-post intervals (Karmarkar et al. 2002; Shouval and Kalantzis 2005). This prediction is based on three observations: calcium influx through NMDARs is a necessary and sufficient signal to induce bidirectional plasticity (Lisman et al. 1998), the sign and magnitude of synaptic plasticity is determined by the calcium concentration in postsynaptic spines (Cormier et al. 2001; Yang et al. 1999), and peak calcium level varies with the time interval between pre- and postsynaptic spiking (Graupner 2010; Karmarkar et al. 2002). These theoretical predictions of pre-before-post tLTD are consistent with experimental evidence in hippocampal slices (Nishiyama et al. 2000; Wittenberg and Wang 2006).

The tLTD at -10 ms and +50 ms flanking the tLTP window at +10 ms and could serve to help sharpen the potentiation of nearly coactive synaptic inputs across the tonotopic map that are generated by the spatio-temporal structure of acoustic stimuli. Frequency and amplitude modulation are common features of natural sounds (Lewicki 2002; Woolley et al. 2005), including species specific vocalizations, and could produce repeated temporal patterns of neural activity in subsets of synapses that could engage STDP as a way of creating either a sensory memory or creating a template within the local circuit based on synaptic strengths for the further analysis of time-varying sounds. The temporally flanking tLTD windows could help to suppress non-coincident synaptic inputs. This idea is consistent with the proposal that recurrent connections can contribute an underlying depolarization that can help to amplify selected afferent signals, and with modeling studies proposing that local amplification could be important in enhancing the sensory selectivity of cortical neurons (Douglas et al. 1995; Krause et al. 2014; Reinhold et al. 2015; Sompolinsky and Shapley 1997).

### Muscarinic modulation of synaptic transmission in AC

Acetylcholine plays an important role in many aspects of cortical development (Hohmann and Berger-Sweeney 1998; Robertson 1998), and some effects of ACh are mediated through activation of mAChRs. Normal cholinergic receptor function appears to be required to help establish the normal tonotopic organization and response features of the auditory cortex. Mice lacking muscarinic M1 receptors more frequently display multi-peak frequency tuning curves as compared to more sharply tuned neurons in wild-type A1. The abnormal tuning curves are also associated with a disorganized tonotopic map (Zhang et al. 2005). Pairing electrical stimulation of nucleus basalis with tones produces large shifts in frequency tuning of A1 neurons (Weinberger 1998) and a corresponding reorganization of the tonotopic map that results in an over-representation of the paired tone frequency (Froemke et al. 2007; Kilgard 1998; Weinberger 1998). However, in M1 receptor knockout mice, pairing nucleus basalis stimulation and tones produces much smaller shifts in frequency tuning in A1 (Zhang et al. 2006). It is not clear, however, to what extent these changes are primarily due to remapping at the level of the thalamocortical recipient cells in layer 4 (whether by thalamic input or by biasing through intracortical circuits), or whether they also reflect changes ascending connections to layer 2/3 or in the layer 2/3 circuitry itself. Tonotopy in A1 is most often measured in anesthetized animals, where the responses are dominated by the more precise tonotopy of layer 4 rather than the imprecise map in layer 2/3 (Kanold et al. 2014; Winkowski and Kanold 2013) (but see (Tischbirek et al. 2019)). Therefore, the specific role of cholinergic systems and M1 receptors in the plasticity of tonotopy or response areas in layer 2/3 cells is not clear.

At the cellular level, acetylcholine acting on mAChRs in cortex can affect intrinsic excitability, synaptic potentials, neurotransmitter release and calcium influx (Cho et al. 2008; Froemke et al. 2007; Metherate and Ashe 1995; Salgado et al. 2007). Consistent with findings in auditory and visual cortices (McCoy and McMahon 2007; Metherate and Ashe 1995) we found that the cholinergic agonists carbachol and Oxo-M depress glutamatergic synaptic transmission. Endogenous activation of mAChRs with an anticholinesterase also produced a weak depression of synaptic potentials suggesting that ambient acetylcholine may tonically regulate synaptic transmission in AC. The synaptic depression generated by carbachol was blocked by atropine, implicating mAChRs rather than nAChRs. The depression of transmission by carbachol was not blocked by 75 nM pirenzepine, which at this concentration is predominantly an M1 receptor antagonist, consistent with results in prefrontal cortex (Vidal and Changeux 1993). In addition, the depression was only weakly antagonized by the M1-selective antagonist VU0255035 (Sheffler et al. 2009) consistent with the suggestion that M1 receptors are not essential in generating the pharmacologically induced synaptic depression.

The muscarinic receptor subtypes M2 and M3 are also expressed in auditory cortex (Salgado et al. 2007). The M2 receptors are localized to excitatory terminals from white matter inputs as well as layer 2/3 GABAergic axon terminals, and presynaptically modulate neurotransmitter release. At the network level, activation of cholinergic synapses may alter the coordinated activity of excitatory and inhibitory neurons by selectively modulating the excitability or synaptic transmission in subtypes of inhibitory cells (Kuchibhotla et al. 2017; Letzkus et al. 2011; Sugihara et al. 2016). On the other hand, our results appear to reveal an effect of mAChR activation with carbachol that is mediated postsynaptically at excitatory synapses onto the pyramidal cells, based on the unchanged paired-pulse ratio of the synaptic responses for both weak EPSPs and for pharmacologically-isolated NMDA receptor currents. Consistent with a postsynaptic site of action, we also found that carbachol increased intrinsic excitability by reducing spike rate adaptation, and enhanced back propagating action potential-mediated calcium influx, likely through reduction in the availability of dendritic potassium conductance. The increase in excitability and dendritic calcium influx are consistent with other studies in auditory and visual cortices (Cho et al. 2008; Metherate and Ashe 1995).

Taken together, these results indicate that activation of mAChRs would increase postsynaptic pyramidal cell excitability while simultaneously decreasing excitatory intracortical transmission. If the effects of mAChRs are selective for intracortical layer 2/3 connections relative to other synaptic inputs, then cholinergic systems could enhance the salience of ascending sensory information arising through thalamocortical afferents (Hsieh et al. 2000) and interlaminar synaptic input from layer 4. A suppression of recurrent excitatory connections with L2/3 by mAChR activation, together with modulation of the inhibitory circuits (Kuchibhotla et al. 2017), might also be expected to alter the frequency sensitivity of superficial AC neurons, by reducing the lateral spread of recurrent excitation.

### mAChR modulation of STDP

There is clear evidence that neuromodulators, including acetylcholine control STDP rules by regulating polarity, magnitude and temporal requirements for plasticity. For example, mAChR activation during pre-before-post pairings has been reported to induce tLTP (Wespatat et al. 2004) and gate tLTD (Seol et al. 2007) in V1. Synaptically-released ACh can enhance tLTP while blocking tLTD in hippocampus (Sugisaki et al. 2011). β-adrenergic receptor activation controls the gating of tLTP in V1 (Seol et al. 2007) and can affect the overall temporal structure of tLTP/tLTD (Salgado et al. 2011). Nicotinic receptor activation prevents tLTP induction in prefrontal cortex (Couey et al. 2007). Dopaminergic activation extends the tLTP window and converts tLTD to tLTP in hippocampus (Zhang et al. 2009). Our results show that mAChR activation with specific agonists or with the anticholinesterase eserine (which may also result in activation of nicotinic receptors) during pre-before-post pairings prevents tLTP induction in AC. The mAChR-mediated suppression of tLTP in AC is consistent with the finding that increasing acetylcholine levels with eserine in CA1, during activation of the cholinergic medial septal inputs can prevent tLTP induction (Sugisaki et al. 2011). The relative timing of glutamatergic versus cholinergic synaptic transmission plays also plays role in the mechanisms and net effects on plasticity (Gu and Yakel 2011). It is not clear how the effects of slow, long-term activation (and potential desensitization) of the receptors, as employed in the experiments here, are related to the effects of the temporally and spatially restricted patterns of cholinergic activity expected in vivo.

The most parsimonious hypothesis to explain the reduction of tLTP in our experiments is that activation of the mAChR’s reduced synaptic transmission during the induction protocol, which in turn resulted in a lower calcium influx that was not sufficient to consistently support tLTP. Three observations support this idea. First, we observed that mAChR activation reduced NMDA current at layer 2/3 synapses, consistent with observations in juvenile rat AC slices (Flores-Hernandez et al. 2009). A molecular mechanism for mAChR-dependent internalization of NMDA receptors has been described in the hippocampus (Jo et al. 2010) that could explain this reduction. Second, tLTP (but not tLTD) was blocked by chelating intracellular calcium with BAPTA, which suggests an obligate role for calcium. This calcium can arise three sources, transmembrane calcium influx through calcium channels opened by depolarization provided by back propagating action potentials, calcium influx through synaptically-activated NMDARs, and calcium-induced calcium release from intracellular stores. Each of these sources likely has different targets because of the spatially-restricted actions of calcium. The increase in calcium influx that we observed in the dendritic shaft during brief trains of action potentials and the increase in the amplitude of that influx during mAChR activation are consistent with similar findings in V1 (Cho et al. 2008). However, it is not clear that these effects directly play a role in regulating tLTP. We did not detect bulk calcium transients in the dendrites associated with our weak (2-8 mV) EPSPs. When EPSPs were paired with an action potential burst, the dendritic calcium transients were larger than with action potentials alone, suggesting an amplification of calcium influx through NMDAR receptors (Kumar et al. 2018; Schiller et al. 1998) by a transient voltage-dependent removal of the Mg2+ block of NMDARs (Nowak et al. 1984). Interestingly, in the presence of carbachol, the dendritic shaft calcium influx during pairing of EPSPs and action potentials did not show enhancement over action potentials alone, consistent with the suppression of NMDA receptor currents by mAChRs (Figure 8). As the activated NMDARs are most likely limited to single dendritic spines and the adjoining dendritic shaft (Müller and Connor 1991) or could expand into more of the shaft area (Eilers et al. 1995) our limited ability to detect small changes in the spines may mean that we missed some key changes in the calcium signal. In addition, if under normal conditions the detected calcium was near the upper end of the non-linear binding relationship between free calcium and the indicator fluorescence, a further increase in calcium may have been masked, so the fluorescence signal did not accurately reflect the intracellular calcium. Although we used two indicators with different reported k_d_ values for these measurements in an attempt to minimize possible effects of binding saturation, the affinity of these indicators for calcium in the cellular environment is not known. These measurements should be revisited with more sensitive and spatially precise methods. Taken together, these results are consistent with the idea that mAChR activation may have reduced the increase in postsynaptic calcium to a level below that required for tLTP induction, most likely as a result of the mAChR-induced reduction in NMDA current.

### Summary

From a functional viewpoint, the depression of tLTP during activation of mAChRs at layer 2/3 synapses suggests that synaptic learning rules can be modified during behavioral states that affect cortical cholinergic tone during the active performance of an auditory task (Kuchibhotla et al. 2017). The reduction of tLTP at the +10 ms interval suggests that a non-modifiable weight may be useful in environmental situations where the attributes of sounds are being identified and tracked, and where synapse-dependent changes and plasticity in sensory processing provided by a normally dynamically changing layer 2/3 circuit in the early parts of the cortical processing pathway would be detrimental to the consistent recognition or discrimination of such sounds. However, these are only one of many distinct synaptic circuits in a complex cortical network, and other synaptic connections may respond to changes in cholinergic tone quite differently.

## Conflict of Interest

The authors have no competing financial interests to declare.

## Author Contributions

D.R. and P.B.M. designed research. D.R. and M.B.K. performed electrophysiological experiments. D.R., M.B.K., and P.B.M. analyzed data and wrote the manuscript.

## Acknowledgements

We thank Dr. Luke Campagnola for his acquisition software, and H. O’Donohue for experimental suggestions on pharmacology and organizational support. This work was supported by US National Institute on Deafness and other Communication Disorders grants R01DC0009809 and R01DC000425 to PBM.

